# Combining hypoxia-activated prodrugs and radiotherapy *in silico*: Impact of treatment scheduling and the intra-tumoural oxygen landscape

**DOI:** 10.1101/856443

**Authors:** Sara Hamis, Mohammad Kohandel, Ludwig J Dubois, Ala Yaromina, Philippe Lambin, Gibin G Powathil

## Abstract

Hypoxia-activated prodrugs (HAPs) present a conceptually elegant approach to not only overcome, but better yet, exploit intra-tumoural hypoxia. Despite being successful *in vitro* and in *vivo*, HAPs are yet to achieve successful results in clinical settings. It has been hypothesised that this lack of clinical success can, in part, be explained by the insufficiently stringent clinical screening selection of determining which tumours are suitable for HAP treatments.

Taking a mathematical modelling approach, we investigate how tumour properties and HAP-radiation scheduling influence treatment outcomes in simulated tumours. The following key results are demonstrated in *silico*: *(i)* HAP and ionising radiation (IR) monotherapies may attack tumours in dissimilar, and complementary, ways. *(ii)* HAP-IR scheduling may impact treatment efficacy. *(iii)* HAPs may function as IR treatment intensifiers. *(iv)* The spatio-temporal intra-tumoural oxygen landscape may impact HAP efficacy. Our *in silico* framework is based on an on-lattice, hybrid, multiscale cellular automaton spanning three spatial dimensions. The mathematical model for tumour spheroid growth is parameterised by multicellular tumour spheroid (MCTS) data.

**Author Summary:** When cancer patients present with solid tumours, the tumours often contain regions that are oxygen-deprived or, in other words, hypoxic. Hypoxic cancer cells are more resistant to conventional anti-cancer therapies, such as chemotherapy and radiotherapy, and therefore tumour hypoxia may complicate treatments. Hypoxia-activated prodrugs constitute a conceptually elegant approach to not only overcome, but better yet, exploit tumour hypoxia. Hypoxia-activated prodrugs are drugs that act as Trojan horses, they are theoretically harmless vehicles that are converted into warheads when they reach their targets: hypoxic tumour cells. Despite being conceptually clever and successful in experimental settings, hypoxia-activated prodrugs are yet to achieve successful results in clinical trials. It has been hypothesised that this lack of clinical success can, in part, be explained by an insufficiently stringent clinical screening selection of determining which tumours are suitable for hypoxia-activated prodrug treatments.

In this article, we investigate how simulated tumours with different oxygen landscapes respond to anti-cancer treatments that include hypoxia-activated prodrugs, either alone or in combination with radiotherapy. Our simulation framework is based on a mathematical model that describes how individual cancer cells in a tumour divide and respond to treatments. We demonstrate that the efficacy of hypoxia-activated prodrugs depends on both the treatment scheduling, and on the oxygen landscape of the specific, simulated tumour.

## Introduction

Oxygen concentrations vary across solid tumours and, although tumours present with high diversity across patients [1], hypoxic regions are prevalent tumour features, commonly provoked by inadequate oxygen supply and high tumour growth rates [2–11]. Hypoxia significantly impacts tumour dynamics, treatment responses and, by extension, clinical outcomes [6,9,12]. Hypoxia may alter cellular expressions of genomes, proteins and epigenetic traits [2], and such hypoxia-induced alterations may cause hypoxic cancer cells to become more resistant to apoptosis [13]. Hypoxia may also alter the metabolism of cells [13], promote angiogenesis by activating associated genes [14] and upregulate efflux systems [15]. Thus hypoxia may both protect and progresses solid tumours [12,13] and, accordingly, severe tumour hypoxia is associated with tumours that are difficult to treat and, by extension, poor prognoses for patients [2,7]. It is well established that hypoxic regions in solid tumours express reduced sensitivity to radiotherapy and a plethora of chemotherapeutic drugs [2,6–9,11,13,14,16–18]. Hypoxic cancer cells in a solid tumour are naturally located far away from active oxygen sources, i.e. blood vessels [7], and therefore drug molecules that are of large size or tightly bound to cell components may not reach hypoxic tumour cells at all [14]. Moreover, genes associated with chemo-resistance may be upregulated by hypoxia [19]. Hypoxia is also regarded to be one of the main factors contributing to radiotherapy failure [14] and radiation-induced DNA damage, especially in the form of double strand breaks, is more easily self-repaired by cells under hypoxic conditions [20].

Due to their severe impact on conventional anti-cancer therapies, such as chemotherapy and radiotherapy, hypoxic cancer cells, and their central mediators [2], have for the last decades been considered to be important treatment-targets [1,14]. In treatment scenarios in which rapid tumour re-oxygenation does not occur, hypoxic tumour regions can, instead, be more directly targeted. In fact, multiple ways to handle tumour hypoxia have been explored. One approach to combating intra-tumoural hypoxia is to increase the tumour oxygenation as part of a neoadjuvant treatment [19]. A second approach to overcome hypoxia is to selectively target hypoxic cancer cells for treatment-sensitising or eradication [4]. A third and conceptually elegant approach to not only overcome, but better yet, exploit intra-tumoural hypoxia is realised by hypoxia-activated prodrugs (HAPs) [14]. HAPs are bioreductive prodrugs that reduce, and thus convert, into cytotoxic agents upon reaching hypoxic (tumour) regions [13,18]. Theoretically, they act as Trojan horses, ideally being essentially harmless until they are converted into warheads in targets, i.e. hypoxic (tumour) regions. The tumour-targeting ability of HAPs is based on the premise that oxygen concentrations in hypoxic tumour regions reach exceptionally low levels, and that such low oxygen levels are much more prevalent in tumours, than in the body tissue that locally surrounds the tumours [13]. Indeed physoxia (the term commonly used to describe oxygen levels found in several types of normal tissue), ranges between 10 and 80 mmHg, and a cancer cell is commonly classified as hypoxic if it has a partial pressure of oxygen (pO_2_) value of 10 mmHg or less [5]. Solid tumours commonly display regions that are even more hypoxic, where pO_2_ values may drop below 5 mmHg [5]. Consequently, HAPs theoretically constitute a means to effectively target hypoxic tumour regions whilst keeping toxic effects localised to tumours, in great part sparing the remaining host system from harmful toxicity and unwanted side effects.

HAPs transform into activated drugs (AHAPs) via reductive metabolism in sufficiently hypoxic environments [3, 14], and the AHAPs can in turn achieve cytotoxic effects in cells [21]. Freely available molecular oxygen may inhibit this bioreduction, and thus HAPs remain (for the most part) more intact, and by extension less toxic, in well-oxygenated environments [13]. Once activated, certain AHAPs may diffuse into their local surroundings. Thus, via bystander effects, for certain HAP drugs, AHAPs may infer damage to cells in which the HAP-to-AHAP bioreduction did not occur. However, a few recent studies dispute the impact of these bystander effects on the overall treatment outcome [22]. In the mathematical model utilised in this study, the dispersion of HAPs and AHAPs obey mechanistic diffusion equations, and the reach of AHAPs can easily be modified by altering coefficients in the AHAP diffusion equation. Thus the influence of bystander effects on the treatment outcome is allowed to range from *negligible* to *highly influential* in our mathematical model.

Multiple HAPs have been evaluated for their clinical potential, both as monotherapies and as part of combination therapies [2,8]. Class I HAPs are activated in moderately hypoxic environments whilst Class II HAPs require more severe hypoxia to undergo the HAP to AHAP bioreduction [23]. One such Class II HAP is evofosfamide, or TH-302, which has been tested in clinical Phase I-III trials [2,19]. TH-302 bioreduces to its activated form, bromo-isophosphoramide mustard (Br-IPM), in hypoxic tumour regions, and Br-IPM is a DNA-crosslinking agent [22]. Multiple *in vitro* and *in vivo* studies have validated this prodrug’s preclincal success and, by extension, its clinical feasibility [6,7,9,10,12,17,21,24–28]. Multimodality treatment strategies combining HAPs, particularly Class II HAPs, with ionising radiation (IR) may be particularly promising [8,9,27–29] as the two therapies conceptually complement each other: HAPs target hypoxic tumour regions whilst radiotherapy is most effective against well-oxygenated tumour regions. Thus, in principal, HAP-IR combination treatments have the ability to produce multifaceted attacks on tumours.

Despite HAPs being conceptually promising and successful in laboratories, their success has not yet been mirrored in clinical trials [1,2,19]. It is hypothesised that this unsuccessful *Bench-to-Bedside* translation is partly due to an insufficiently stringent clinical screening practice of selecting tumours that are suitable for HAP treatments [19]. It is likely that some of the tumours enrolled in clinical trials have been insufficiently hypoxic to benefit from treatment plans involving HAPs [1]. To investigate this hypothesis, we here propose a mathematical modelling angle to simulate how spatio-temporal tumour features may impact HAP efficacy and how scheduling influences the outcome of multimodality HAP-IR treatments *in silico*.

Today, mathematical modelling constitutes an indispensable complement to traditional cancer research [30]. Models provide an opportunity to study biological phenomena *in silico* that may not be empirically observable and, moreover, *in silico* experiments are fast and cheap to run, easy to reproduce and not directly associated with any ethical concerns. Previous mathematical studies have already contributed to the overall understanding of HAPs, quantified key mechanisms associated to them and illustrated their clinical feasibility. Foehrenbacher *et al.* [31] deployed a Green’s function method, in customised form, and pharmacokinetic/pharmacodynamic (PK/PD) modelling to quantify anti-cancer bystander effects elicited by the HAP PR-104 in a simulated, three-dimensional tumour comprising a microvascular network. Another concurrent article used similar mathematical concepts to compare Class I HAPs to Class II HAPs and, furthermore, to determine optimal properties for Class II HAPs [23]. Lindsay *et al.* [32] developed a stochastic model to study monotherapies and combination therapies involving HAPs, specifically TH-302, and erlotinib. Amongst other findings, they concluded that a combination therapy of the two drugs impedes the uprising of drug resistance. Since HAPs bioreduce to their activated form under hypoxic conditions it follows that AHAP activity increases with intra-tumoural hypoxia. Accordingly, a previous study by Wojtkowiak *et al.* [33] conceptually validated the strategy of amplifying TH-302 activity by deliberately exacerbating intra-tumoural hypoxia using exogenous pyruvate. Their study combined mathematical modelling with metabolic profiling and EPR (electron paramagnetic resonance) imaging. HAP dynamics were modelled using reaction-diffusion/convection equations coupled with fluid-structure interactions. In line with these previous mathematical studies, the aim of this *in silico* study is to contribute HAP-related insights gained by mathematical modelling, according to a *Blackboard-to-Bedside* [34] approach.

## Model

An on-lattice, hybrid, multiscale cellular automaton (CA) is here used to model solid tumours subjected to HAP and IR monotherapies, as well as HAP-IR combination therapies. Tumour growth and HAP responses are parameterised by published data from an *in vitro* study performed by Voissiere *et al.* [35], in which multicellular tumour spheroids (MCTSs) where grown and exposed to HAPs. Specifically, we use their data for human chondrosarcoma HEMC-SS cells exposed to the hypoxia-activated prodrug TH-302. Our mathematical model is thereafter extended to simulate *in vivo* drug dynamics in order to investigate scheduling aspects of HAP-IR combination therapies. The parameters used in this paper can be modified in order to simulate specific cell-lines and drugs, and model rules can be altered in order to simulate both *in vitro* and *in vivo* cancer cell populations, MCTSs or tumours. Thus, upon the availability of appropriate data, various tumour scenarios and treatment schedules and doses can be investigated *in silico*. Hence the mathematical model presented here constitutes a valuable and versatile complement to both *in vitro* and *in vivo* experiments. The model used in this study is an extension of a previous, well-established model presented by Powathil *et al.* [36]. All parameters used in the model are motivated from experiments and literature, as described throughout this section, and are summarised Table 1.

**Table 1.** A summary of model parameters used in the mathematical framework.

| Equation | Parameter | Value |
| --- | --- | --- |
| Cellular Automaton |  |  |
| N/A | $\Delta x_1 = \Delta x_2 = \Delta x_3$ (spacing)<br>$\Delta t$ | 20 $\mu\text{m}$<br>0.001 hours |
| Cell-cycle and proliferation |  |  |
| N/A | $\mu, \sigma$<br>$\theta_{G1}, \theta_S, \theta_{G2}, \theta_M$<br>$a_1, a_2, a_3$<br>$\nu$ | 40 hours, 4 hours<br>$\frac{11}{24}, \frac{8}{24}, \frac{4}{24}, \frac{1}{24}$<br>0.9209, 0.8200, -0.2389<br>3 |
| Oxygen |  |  |
| 5 | $D_K(x, t) = \begin{cases} D_K/1.5 & \text{if cell in } (x, t) \\ D_K & \text{otherwise} \end{cases}$<br>$cell(x, t) = \begin{cases} 1 & \text{if viable cell in } (x, t) \\ 0 & \text{otherwise} \end{cases}$<br>$m(x, t) = \begin{cases} 1 & \text{if } (x, t) \text{ outside MCTS} \\ 0 & \text{otherwise} \end{cases}$ | $D_K = 2.5 \times 10^{-5} \text{ cm}^2\text{s}^{-1}$ |
| 6 | h | 0.5 mmHg |
| 11 | OER<sub>m</sub> | 3 |
| 12 | K<sub>m</sub> | 3 mmHg |
| Drugs |  |  |
| 7 | b | (hour)<sup>-1</sup> |
| 8 | [pO<sub>2</sub>]<sub>50</sub> | 0.2 mmHg |
| 9 and 10 | $D_{HAP}, D_{AHAP}$<br>$\eta_{HAP}, \eta_{AHAP}$ | $2 \times D_K(x, t), \frac{1}{4} \times D_K(x, t)$<br>*picked from half-life times:*<br>$t_{1/2, HAP}=0.81$ hours,<br>$t_{1/2, AHAP}=0.70$ hours |
| 9 | $T_{L \rightarrow R}$ (for the no removal *in vitro* case) | Infinity |
| Radiotherapy |  |  |
| 13 | $\alpha(G1), \beta(G1)$<br>$\alpha(S), \beta(S)$<br>$\alpha(G2), \beta(G2)$<br>$\alpha(M), \beta(M)$<br>$\alpha(G0), \beta(G0)$ | 0.351, 0.04<br>0.1235, 0.04<br>0.793, 0<br>0.793, 0<br>$\alpha(G1)/1.5, \beta(G1)/(1.5^2)$ |

### Mathematical Framework: A Cellular Automaton (CA)

The CA used in this model allows for spatio-temporal dynamics and intra-tumoural heterogeneity including variations in cell-cycle progressions, oxygen levels, drug concentrations and treatment responses amongst cancer cells [34, 36–38]. The model is multiscale and integrates both intracellular and extracellular regulations. *In vitro* experiments have demonstrated that MCTSs are more HAP-sensitive than are monolayers. This increase in sensitivity has been attributed to the microenvironment correlated to multilayer cultures [17]. Aspiring to achieve an *in silico* model that is as clinically relevant as possible, we here let the CA lattice extend in three spatial dimensions. The lattice is specifically a square lattice containing 100^3^ lattice points, simulating a physical environment of (2mm)^3^. Thus each voxel in the lattice spans a volume of (20*μ*m)^3^ and each lattice point may be occupied by either one cancer cell or extracellular space. These dimensions agree with previous mathematical studies [36], and reported cell population densities in the MCTSs that are used to calibrate the model [35]. The time step used for the temporal progression of the CA is Δ*t* = 10^-3^ hours, by appropriate non-dimensionalisation of oxygen dynamics [36].

### Cell-Cycle Progression

On an intracellular scale, sub-cellular mechanisms are modelled individually for each cell in order to allow for variations amongst cancer cells. Cell-cycle progression is one such intracellular process, and it is here governed by an intrinsic cell-cycle clock attributed to each individual cell. In order to account for cell-cycle asynchronicity amongst cells, each cell *i* is assigned an individual, stochastic doubling-time *τ_i_* which corresponds to the time it takes for a cell to complete one cell-cycle, and double by producing a daughter cell, under well-oxygenated conditions. Here, *τ_i_* is picked from a normal distribution [37] with a mean value *μ* and a standard deviation *σ*, which are picked to match cell population growth-rates reported from Voissiere *et al.* [35], as demonstrated in Figure 1.

**Fig 1.**
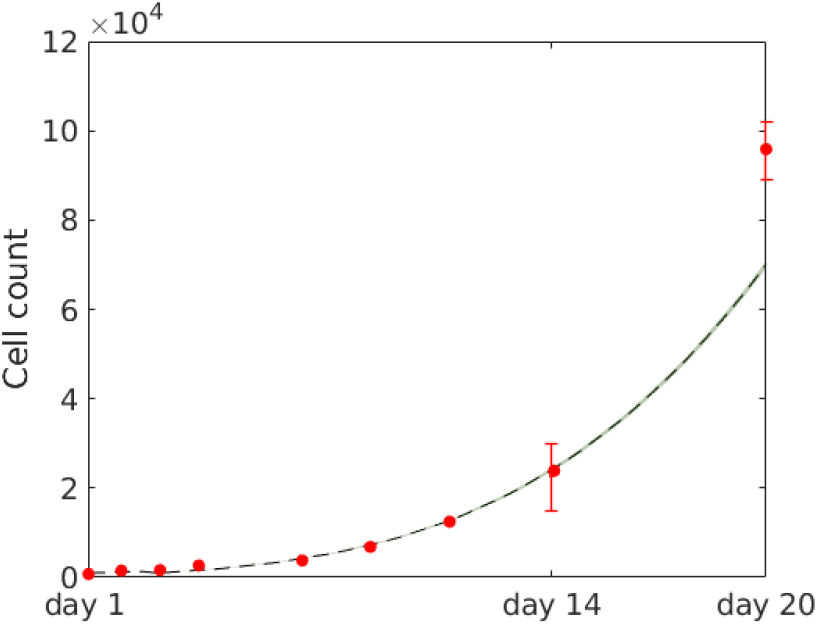
Cell count over time for tumour spheroids. The *in silico* data is based on 10 simulations runs, the mean (black line) shows the average value and the gray ribbon shows standard deviation. *In vitro* data (red error bars) are extracted from plots produced by Voissiere *et al.* [35] using a Java program (DataThief III [39]).

As sensitivity to radiotherapy is cell-cycle dependent [20], it is important to track cell-cycle phase progression in the model. Thus each cell in the model follows a cell-cycle typical to that of eukaryotic cells and, in particular, a cell is defined to be in the gap 1 (G1), synthesis (S), gap 2 (G2) or mitosis (M) phase of the cell-cycle. Under well-oxygenated conditions, the fraction of time spent in each of the four distinct cell-cycle phases are Θ_*G*1_, Θ_*S*_, Θ_*G*2_ and Θ_*M*_ for the cell-cycle phases G1, S, G2, M respectively, where the Θ-fractions sum up to one so that

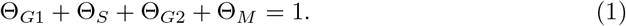

The four theta values are picked from literature in order to match typical cell-cycle phase lengths of rapidly cycling human cells with a doubling time of roughly 24 hours [40]. Specifically, we set the G1, S, G2 and M phase to respectively occupy 11/24:ths, 8/24:ths, 4/24:ths and 1/24:th of a cell’s individual doubling time. These values can be amended upon availability of cell-line specific data. Thus the time spent in each of the four distinct cell-cycle phases, for a well-oxygenated cell *i* with a doubling time *τ_i_*, is here Θ_*G*1_ *τ_i_*, Θ_*S*_*τ_i_*, Θ_*G*2_*τ_i_* and Θ_*M*_*τ_i_* for the cell-cycle phases G1, S, G2 and M respectively so that

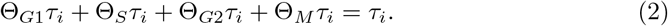

However, low cellular oxygen levels have been shown to delay cell-cycle progression by inducing arrest in particularly the G1 phase of the cell-cycle [41]. Mathematically, the cell-cycle can be modelled in various ways. For example, in mechanistic cell-cycle models derived by Tyson and Novak [42], the cell-cycle is governed by a regulatory molecular network that can be described by a system of ordinary differential equations. By incorporating hypoxia-induced factors in the system of equations, the G1 phase can be inherently elongated under hypoxic conditions [36]. In this study, however, cell-cycle progression is merely modelled using a phenomenological clock, instead of a more detailed Tyson-Novak type of model. As a result of this, there is no mechanistic functionality driving G1-arrest under hypoxic conditions in our model. To remedy this fact, we here introduce an additional function to achieve an oxygen-dependent elongation of the G1-phase. We name this function the G1 Delay Factor (*G*1*DF*) such that,

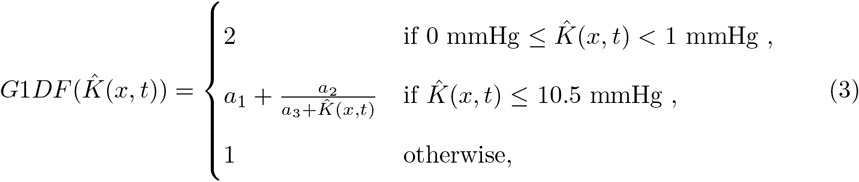

where 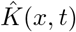 denotes the oxygenation (in units of mmHg) of a cell in point *x* at time *t*. The G1DF, which is illustrated in Figure 2, is an approximation for how much the G1 phase is expanded in time as a function of oxygen pressure, here measured in units of mmHg. The G1DF is matched to fit data points extracted from a previous mathematical study by Alarcon *et al.* [41], in which a Tyson-Novak cell-cycle model is extended to incorporate the action of p27, a protein that is upregulated under hypoxia and delays cell-cycle progression. Thus here the time spent in the *G*_1_ phase, *τ*_*G*1_, is given by

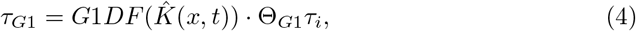

where 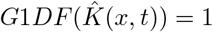 for normoxic cells. The lengths of other cell-cycle phases are approximated as non-oxygen dependent in the model.

**Fig 2.**
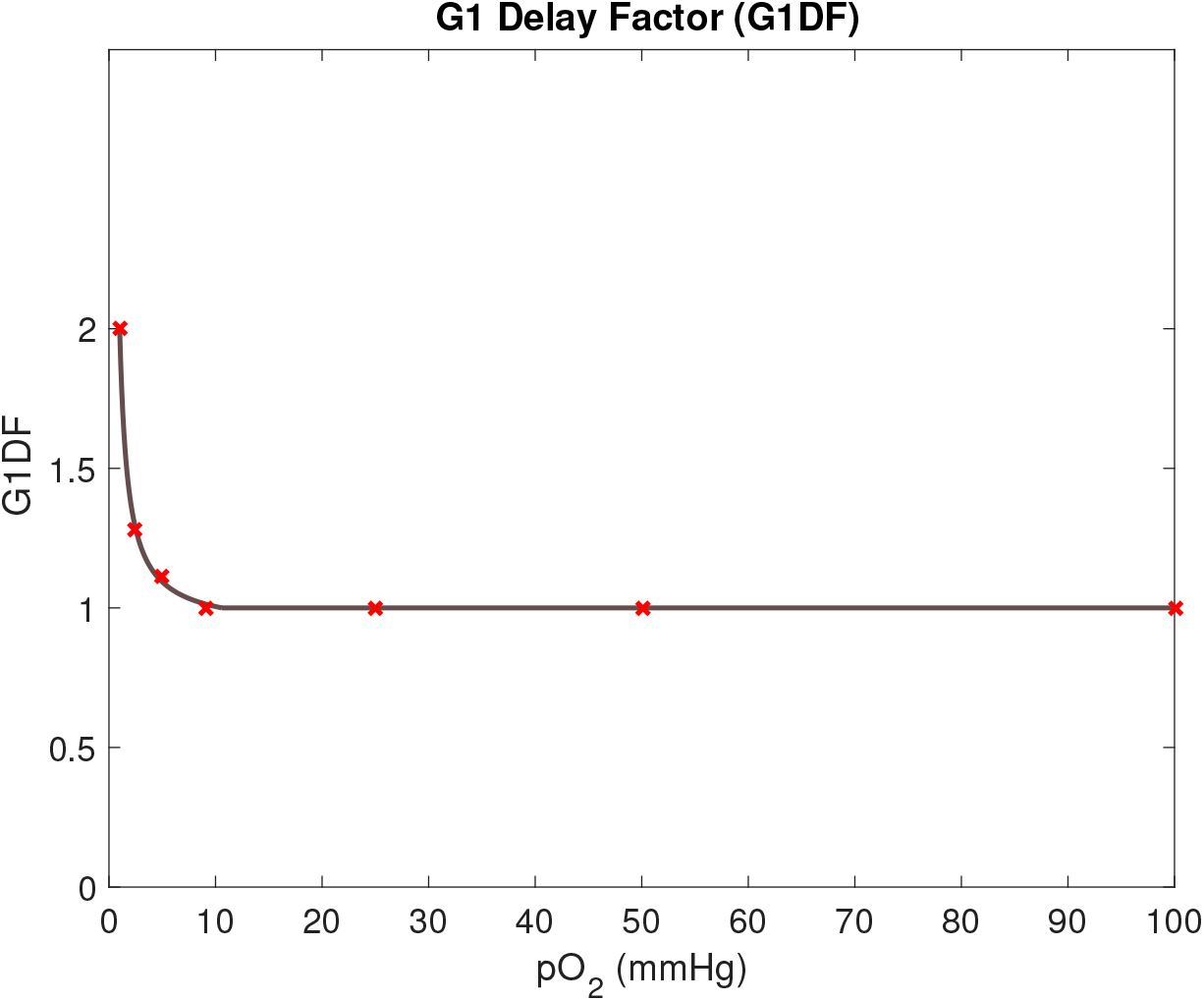
The G1 Delay Factor (G1DF) is incorporated in the model to achieve oxygen-dependent G1 arrest. The G1DF (dark line) is extrapolated from *in vitro* data (red crosses) from a previous mathematical study by Alarcon *et al.* [41].

### Tumour Growth

In the model, a tumour is grown from one seeding cancer cell which divides and gives rise to a heterogeneous MCTS. Each cell that is placed on the lattice commences its first cell-cycle in the G1 phase and once a viable, i.e. undamaged, cell has completed the mitosis (M) phase of the cell-cycle, a secondary cell, namely a daughter cell, is produced and placed in the neighbourhood of its mother cell. In the model, cell-division occurs provided that free space is available on the lattice in the *ν*th order neighbourhood of the mother cell, where the value for *ν* is fitted form experimental data [35]. This constraint simulates a scenario in which cell-division is inhibited by some lack of resources such as space or nutrients. (By setting *ν* = ∞, the model can be adapted to disregard these spatial cell-division constraints [37]). If no free space is available in the vth order neighbourhood of a mother cell that is ready to divide, no daughter cell is produced, and instead the mother cell assumes a state in which it progresses through the cell-cycle very slowly (simulating an *in vitro* spheroid case, in which inner cancer cells experimentally have shown a reduced proliferation rate [35]), or not at all (simulating an *in vivo* case in which cells may enter a quiescent G0 phase [36]). Should neighbourhood space be made available again, as a result of cells getting removed from the lattice in response to anti-cancer treatments, such slow-cycling or resting cells may re-assume an actively cycling state. When cell-division occurs, a daughter cell is placed on a random lattice point in the neighbourhood of the mother cell, where up to *ν* spherical neighbourhoods are regarded and lower order neighbourhood are occupied first. To accomplish spherical-like tumour growth the model stochastically alternates between deploying Moore and von Neumann neighbourhoods [36] for daughter cell-placements. In order to agree with the MCTS data [35] used to calibrate the model, we here pick *ν* = 3, as illustrated in Figure 3, and thus a daughter cell may be placed up to three neighbourhoods away from its mother cell. Note that, in the work presented by this paper, neither necrotic nor apoptotic tumour cells are included in the pre-treatment tumour growth model, and instead we make the simplifying modelling assumption that the density of viable cells is constant (one cancer cell per lattice point) within the simulated MCTSs before any treatment is given. However, CA are easily adaptable and thus, if appropriate, modelling rules concerning necrotic and/or apoptotic cells can be included in the mathematical framework. The *in vitro* experiment produced and reported by Voissiere *et al.* [35] does detect apoptotic cells in the MCTSs, where these are primarily located towards the center of the spheroids.

**Fig 3.**
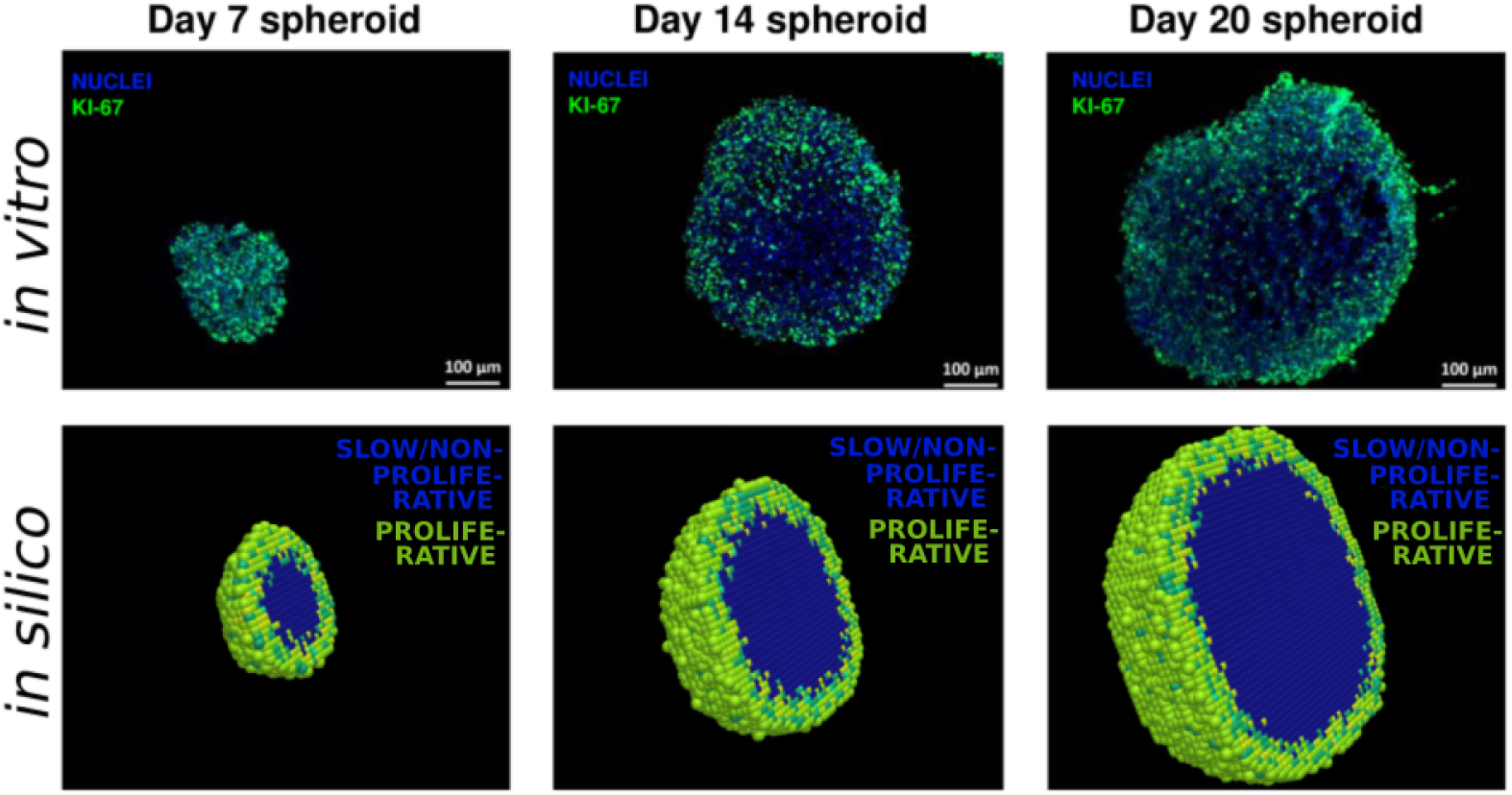
Images from *in vitro* experiments performed by Voissiere *et al.* [35], in which cell nuclei are stained blue and proliferative cells are stained green by the proliferation marker Ki-67. Bottom: Images from *in silico* experiments performed in this study, where proliferative (cycling) cells are coloured green and inner (slow-cycling or non-proliferative) cells are coloured blue.

### Oxygen Distribution and Hypoxia

Oxygen is assumed to be readily available outside the tumour and, therefore, lattice points outside the tumour are oxygen source points in the model. Viable (i.e. non-damaged) cells are modelled as oxygen sinks as they consume oxygen in order to function. The distribution of oxygen across the lattice is modelled by a mechanistic partial differential equation (PDE), specifically a reaction-diffusion equation such that

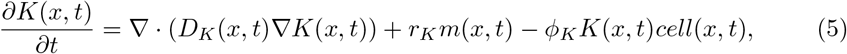

coupled with no-flux boundary conditions. Here *K*(*x, t*) denotes the oxygen level in lattice point *x* at time *t*. *D_K_* (*x, t*) is the diffusion coefficient, which is higher in lattice points occupied by cells compared to unoccupied lattice points, so that oxygen diffuses slower over cancer cells than in extracellular space in the model [36]. The binary function *cell*(*x,t*) is equal to one if the lattice point is occupied by a viable cancer cell, and zero otherwise. Similarly, the binary function *m*(*x, t*) is one if the lattice point is outside the tumour and zero otherwise, i.e. *m*(*x,t*) = 1 if the lattice point in (*x,t*) is neither occupied, nor enclosed, by cancer cell(s). The oxygen production rate is denoted by *r_K_* and the cellular oxygen consumption rate is *ϕ_K_*. Thus the first term in the Eq 5 describes oxygen diffusion, the second term describes oxygen sources and the final term describes cellular oxygen consumption. In the model, the diffusion coefficient for oxygen is gathered from literature but the production and consumption rates are calibrated *in silico* to match *in vitro* data from Voissiere *et al.* [35], specifically to achieve appropriate oxygen gradients. Note that the no-flux boundary condition causes the total amount of oxygen on the lattice to increase over time. To express oxygenation levels on the lattice in scaled form, a scaled oxygen variable 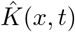 is introduced which is obtained by

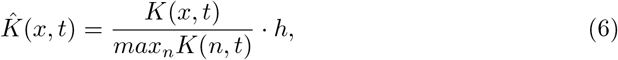

where *max_n_K*(*n, t*) denotes the maximal *K*(*x,t*)-value (of all *n* lattice points) at time *t* [43]. The scaling-factor, *h*, (with unit mmHg), is incorporated in order to calibrate the model to fit MCTS data [35], as illustrated in Figure 4. Note that an alternative way of incorporating oxygen distribution in the model (without having to re-scale the oxygen concentration) is by using an oxygen source term that is proportional to the difference between some reference oxygen concentration *K_υ_* (measured inside oxygen sources e.g. vessels) and the oxygen concentration in the rest of the domain. In the model, a cell is defined to be hypoxic if it has a scaled oxygen value such that 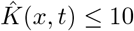 mmHg [36] and the 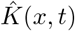 influences G1-arrest (Figure 2), radio-sensitivity (Figure 7) and HAP-AHAP bioreduction rates (Figure 5).

**Fig 4.**
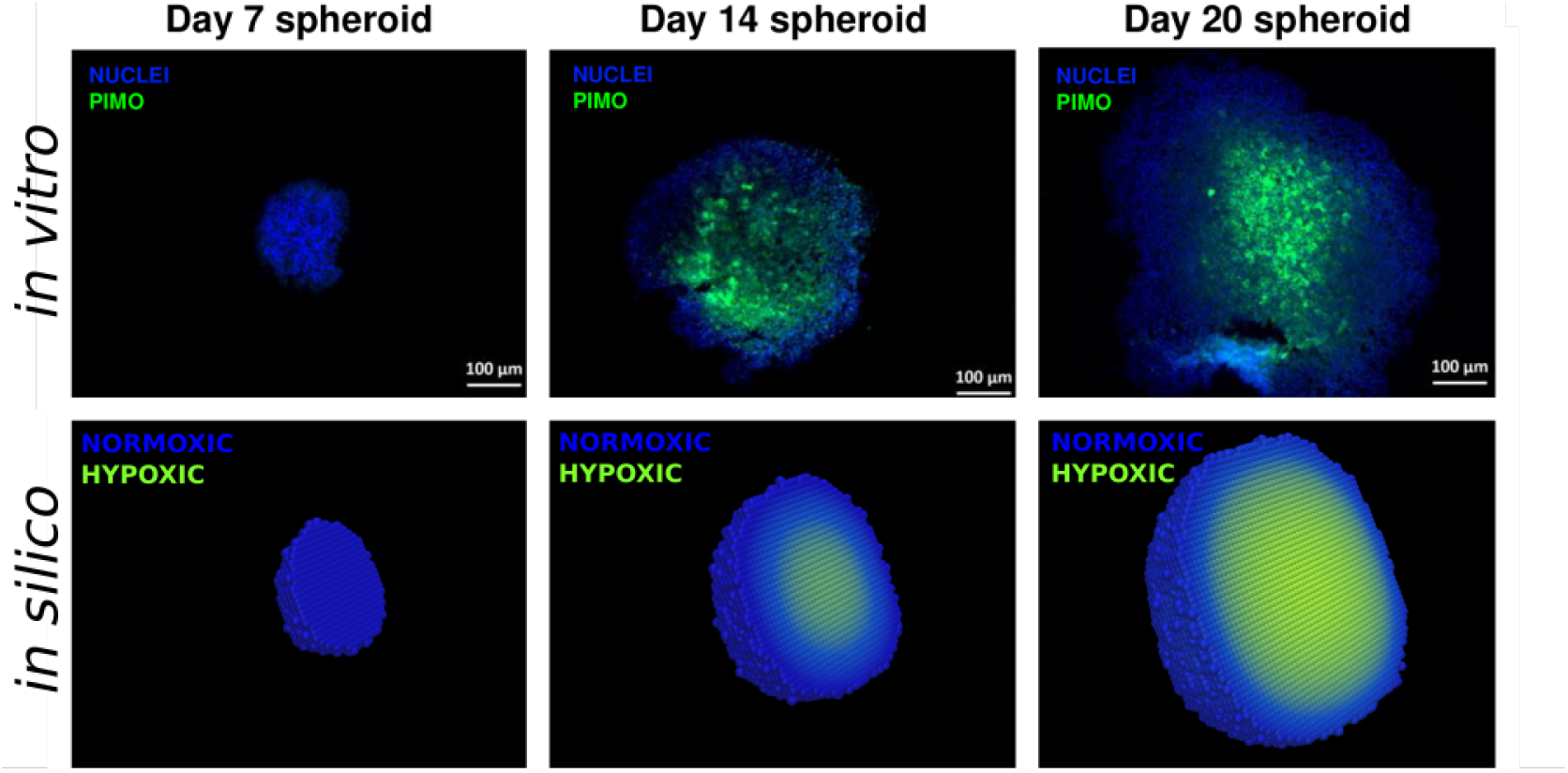
Top: Images from *in vitro* experiments performed by Voissiere *et al.* [35], in which hypoxic cells are stained green by pimonidazole and normoxic cells are stained blue. Bottom: Images from *in silico* experiments performed in this study, where hypoxic cells (*pO*_2_ ≤ 10 mmHg) are coloured green and normoxic cells (*pO*_2_ > 10 mmHg) are coloured blue.

**Fig 5.**
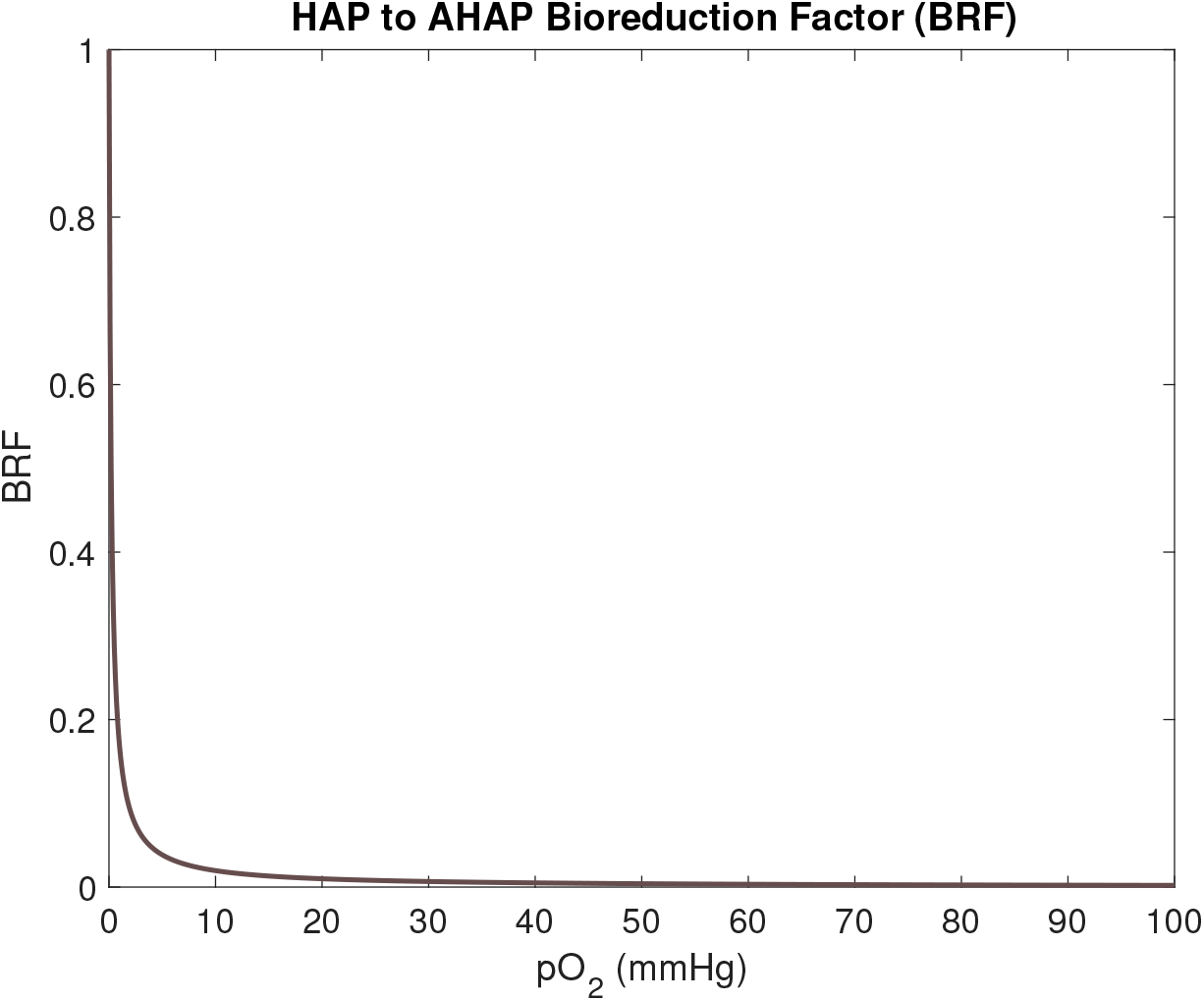
The bioreduction factor, *BRF*, expresses the fraction of HAP compound that reduces to AHAP compound within one hour as a function of oxygenation (in mmHg).

### Hypoxia-Activated Prodrugs

Anti-cancer prodrugs constitute relatively harmless compounds in their inactivated form with the potential to bioreduce, or transform, into cytotoxic species [21]. Specifically for HAPs, this bioreduction occurs in hypoxic conditions and thus HAPs are able to selectively target hypoxic tumour regions [21]. The oxygen dependent bioreduction is here modelled by the function 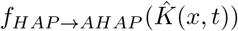, where

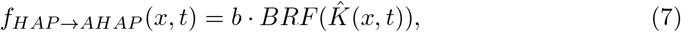

where *b* is a time-scaling factor with and *BRF* is a bioreduction factor as illustrated in Figure 5 and

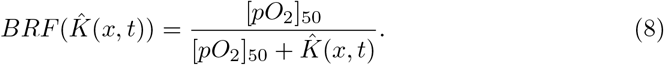

Here [*pO*_2_]_50_ denotes the oxygen value yielding 50% bioreduction (in one hour), chosen to be 0.2 mmHg, for evofosfamide, as is done in a previous mathematical model by Hong *et al.* [44]. As illustrated in Figure 5, the BRF value rapidly decreases for *pO*_2_ values (i.e. 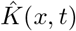 values) between 0 and 10 mmHg.

The mechanistic reaction-diffusion equations governing the distribution of HAPs and AHAPs across the lattice are respectively given by [45]

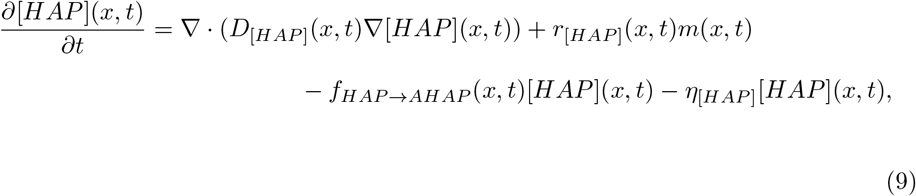

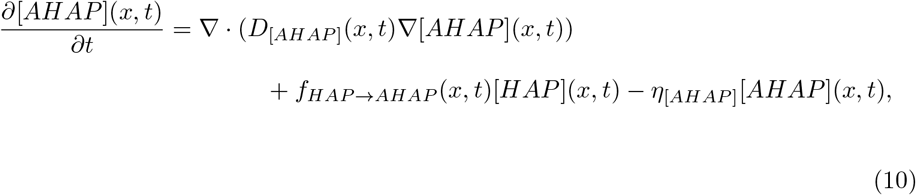

where [*HAP*](*x,t*) denotes the concentration of HAPs and [*AHAP*](*x,t*) denotes the concentration of AHAPs in point *x* at time *t*. *D*_[*HAP*]_ (*x,t*) and *D*_[*AHAP*]_ (*x,t*) denote the respective diffusion coefficients, *r*_[*HAP*]_ (*x,t*) denotes the HAP production rate, *η*_[*HAP*]_ and *η*_[*AHAP*]_ denote the corresponding decay rates. AHAPs are harmful agents which are here assumed to inflict damage that is cell-cycle non-specific. Consequently, cells that are in any cell-cycle phase (G1, S, G2, M), including cells that are in a slow or non-cycling state in the centre of the MCTS, are susceptible to AHAP-inflicted damage in the model. A cell in point *x* at time *t* is damaged by the cytotoxic AHAPs if [*AHAP*](*x, t*) ≥ Ψ, where Ψ is the lethal AHAP concentration threshold. Note that using a threshold value to determine cell fate (death) is a model approximation and, in reality (*in vitro/in vivo*), cellular drug responses will depend on several drug (pharmacokinetic/pharmacodynamic) factors, as well as specific cell/tumour details. When a cell dies, it reduces to a membrane-enclosed cell-corpse which is (in *vivo*) digested by macrophages [46]. In the model, the time it takes between a cell is declared dying and it is removed from the lattice is denoted *T_L→R_* (L for lethal event, R for removal). Three cases for this time span *T_L→r_* are investigated in this study: (*i*) the first extreme case in which a dead cell is *never* removed from the lattice (simulating an *in vitro environment*), (*ii*) the other extreme case in which a cell is *instantaneously* removed from the lattice upon receiving lethal damage, and (*iii*) a mid-way case in which a cell is removed from the lattice after a time-period corresponding to its doubling time has passed, i.e. *T_L→R,i_* = *τ_i_*. Results using the first case are included in the main text of this manuscript, and results for cases (*ii*) and (*iii*) are provided in the supplementary material in which we demonstrate that, within the scope of the performed *in silico* experiments, this choice of *T_L→R_* value does not affect our qualitative findings.

#### Parameters

In our mathematical model, extracellular space (i.e. lattice points outside the tumour) are HAP source points, and from there HAPs are quickly distributed across the lattice. Drug transportation of HAPs from source points to cells is mediated only by the diffusion terms in Eq 9 and similarly AHAP transportation is mediated only by the diffusion term in Eq 10. Consequently, the drug diffusion coefficients *D*_[*HAP*]_ and *D*_[*AHAP*]_ represent all biophysical drug transportation across the lattice *in silico*. HAPs must possess certain appropriate attributes in order to produce desired effects [17]. For example, HAPs should be able to travel relatively long distances without being metabolised, specifically distances longer than that of which oxygen travels, in order to reach hypoxic tumour regions. As oxygen is consumed by the cells, whilst HAPs require certain micro-environmental conditions to be met in order to metabolise, HAPs may reach regions located relatively far away from blood vessels, that oxygen can not reach. It has, indeed, been demonstrated *in vivo* that TH-302 has the ability to reach hypoxic regions, where it is activated [47]. Conversely, AHAPs should ideally travel relatively short distances in order to localise AHAP activity to tumour regions only, and thus to minimise unwanted extra-tumoural toxicity. The diffusion length of oxygen is reported in literature to be approximately 100 *μ*m [36] however, to our knowledge, no diffusion length of neither TH-302 nor Br-IPM has been explicitly reported. However, the diffusion length of the HAP/AHAP pair AQ4N/AQ4 has been shown to be reach roughly 1.5 times that of oxygen (or 150*μ*m) in xenografts [48]. With this motivation, we here approximate the diffusion coefficient of TH-302 to be twice that of oxygen. (*This according to the relationship* 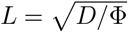, *where L is the diffusion length scale*, Φ *is the compound uptake and the diffusion coefficient of a certain compound, D, is proportional to L*^2^, *neglecting details of compound uptake [36]. Thus here we make the simplified approximation that* 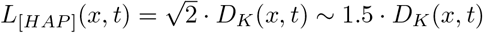.) Similar to previous procedure, the diffusion length of AHAPs is approximated to be half that of oxygen from which it follows that *D*_[*AHAP*]_(*x,t*) = (1/4) · *D_K_*(*x,t*). These parameter estimations suffice to conceptually, and qualitatively, describe the nature of HAPs and AHAPs, but can be amended upon the availability of new data. By adjusting the diffusion coefficients *D*_[*HAP*]_ and *D*_[*AHAP*]_, the influence of bystander effects are allowed to range from negligible to highly influential in our mathematical framework.

The half-life times of TH-302 and Br-IPM have been reported to be 0.81h and 0.70h respectively in a clinical trial [11], these values are used to determine the decay rates *η*_[*HAP*]_ and *η*_[*AHAP*]_. This half-life time of TH-302 is in accordance with preclinical predictions obtained from allometric scaling [26]. Note that the drug decay coefficients, *η*_[*HAP*]_ and *η*_[*AHAP*]_ in Eq 9 and Eq 10 respectively, simulate all drug clearance from the system, i.e. both metabolic clearance and excretion.

### Radiotherapy

Cellular responses to radiotherapy are dependent on oxygenation status [4], cell-cycle phase [49,50], and cell-line characteristics. Cellular radiotherapy responses are here modelled using an appropriate CA adaptation of the widely accepted Linear-Quadratic (LQ) model. In the traditional LQ model, the survival fraction of a cell population is given by *S*(*d*) = *e*^−*nd*(*α+βd*)^, where *d* is the radiation dosage, *n* is the number of administered radiation fractions and *α* and *β* are cell-line specific sensitivity parameters [51]. In order to include cell-cycle sensitivity, *α* and *β* are here cell-cycle dependent and the oxygen modification factor (OMF) is incorporated to include oxygen sensitivity [52], such that

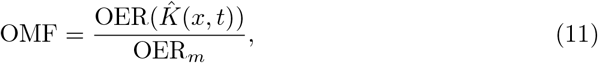

where

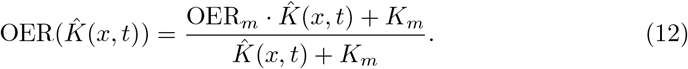

Here, OER_*m*_ = 3 is the maximum value under well-oxygenated conditions and *K_m_* = 3 mmHg is the *pO*_2_ value achieving half of the maximum ratio [43]. The OER and OMF functions are illustrated in Figure 6.

**Fig 6.**
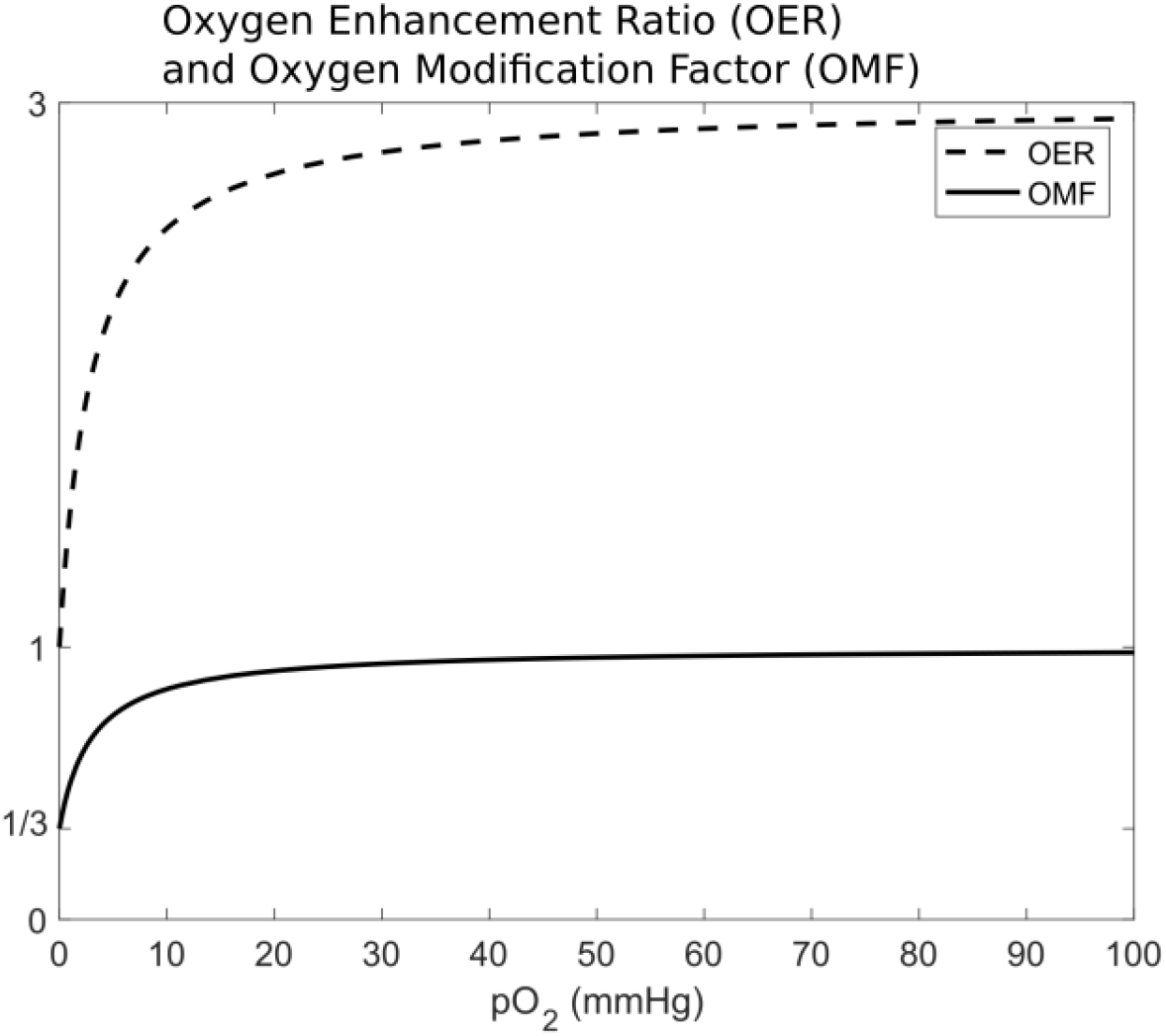
The Oxygen Enhancement Ratio (OER) and the Oxygen Modification Factor (OMF) are incorporated in the mathematical model to quantify the influence of oxygen on radiotherapy responses. Cells are the least radiosensitive for low pO_2_ values. The OER and OMF curves have steep gradients between the oxygen values 0 and 10 mmHg, after which they respectively asymptote to the values 3 and 1 for higher oxygen values.

The survival probability of a cell in point *x* at time *t* is here given by

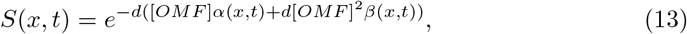

where the cell-cycle phase specific *α* and *β* values are gathered from a previous study by Kempf *et al.* [53], and are listed in Table 1. Cellular responses to a 2Gy IR dose for a generic cancer cell-line, as a function of oxygenation and cell-cycle phase details, are illustrated in Figure 7.

**Fig 7.**
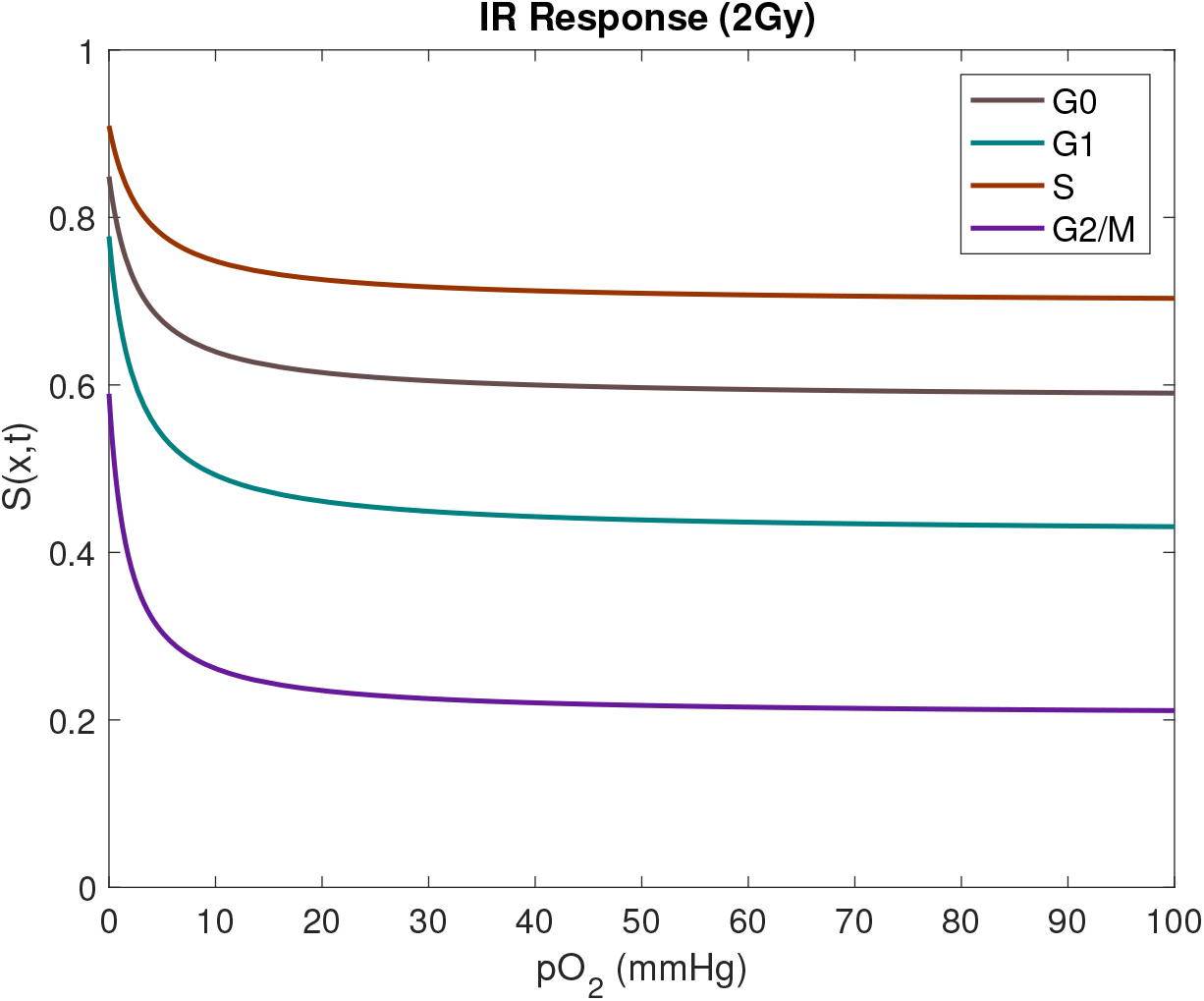
The probability that a cell, in the mathematical model, exposed to a radiotherapy dose of 2 Gy survives. The survival probability *S*(*x,t*) is function of a cell’s current cell-cycle phase and oxygenation value. Cells are the most likely to survive radiotherapy when hypoxic.

### Parameters

In this study we attempt to replicate the nature of generic eukaryotic cell-lines, the HAP evofosfamide (TH-302) and its corresponding AHAP, Br-IPM. The parameters, which are listed in Table 1, are chosen accordingly but can be adapted to represent other specific cell-lines or drugs upon data becoming readily available.

### Implementation and *in silico* Framework

The mathematical model is implemented in an in-house computational framework written in C++ deploying high-performance computing techniques. The PDEs describing oxygen and drug distribution across the lattice are solved using explicit finite difference methods with no-flux boundary conditions. Maps of cancer cells and the microenvironment are visualised in ParaView [54]. Using this computational framework, various experimental *in vitro* and *in vivo* scenarios are formulated and simulated *in silico*. In order to grow an *in silico* MCTS, one seeding cancer cell is placed on the lattice, this cell divides and gives rise to a MCTS that is heterogeneous in nature, as in-built model stochasticity creates cell-cycle asyncronosity amongst tumour cells [55], and oxygen levels vary across the MCTS. Such virtual spheroids are thereafter subjected to various treatments comprising HAPs and/or IR. Treatment commence when MCTSs consist of, in the order of, 100,000 cancer cells or ‘agents’ in our agent-based model. Due to the high number of agents, and the fact that the intrinsic model stochasticity only involves a few events during the simulated treatment time (specifically 0-3 cell divisions and potentially one response to radiotherapy) the quantitative results do not differ much between *in silico* runs. Performing the same *in silico* experiment 10 times yields a standard deviation that can be regarded as negligible (as the standard deviations obtained in this study are less than 0.5% of the mean values). From this we argue that basing our results on mean values from 10 simulation runs per experiment is enough to mitigate intrinsic model stochasticity to a sufficient level for this qualitative study.

## Results and Discussion

In the following sections, we compare treatment responses in two different *in silico* tumour spheroids, specifically a ‘Large’ and more hypoxic MCTS and a ‘Small’, less hypoxic MCTS. The ‘Small’ MCTS corresponds to the 20 day-old MCTS in Figures 3 and 4, that is calibrated by *in vitro* data from Voissiere *et al.* [35]. The ‘Large’ MCTS is extrapolated by letting the ‘Small’ MCTS grow for yet another 10 days *in silico*, until it reaches an age of 30 days. The ‘Small’ and ‘Large’ MCTSs are illustrated in Figure 8.

**Fig 8.**
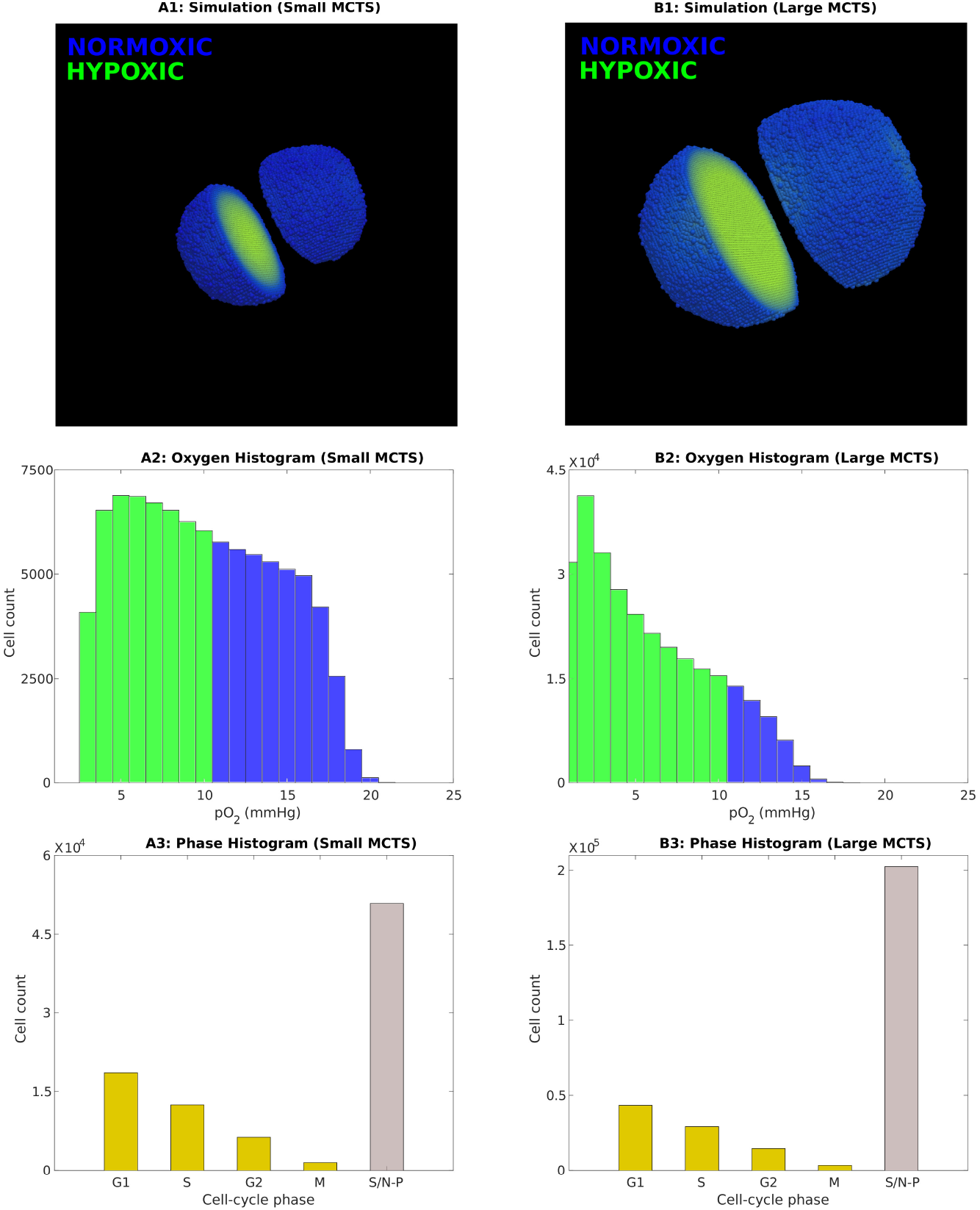
The ‘Small’ (20 day old) MCTS and the ‘Large’ (30 day old) MCTS are used to allow for comparisons in treatment responses between tumours with different oxygenation levels. Top: Simulation snapshots of the MCTSs at the time point *T*_0_ when treatments commence (A1: Small MCTS, B1: Large MCTS). Hypoxic cells (pO_2_ ≤ 10 mmHg) are green whilst normoxic cells are blue. Middle: Oxygen histograms at time *T*_0_, in which hypoxic cell counts are shown in green and normoxic cell counts are shown in blue (A2: Small MCTS, B2: Large MCTS). Bottom: Cell-cycle phase histograms at time *T*_0_ (A3: Small MCTS, B3: Large MCTS). The slow/non-proliferative, inner cancer cells are labeled S/N-P.

The simulated IR dose is chosen to be 2 Gy, and to allow for intuitive comparisons between the two different monotherapies, the HAP dose (Dose_*HAP*_) is here qualitatively chosen, and calibrated to yield the a similar *in silico* response as the 2 Gy IR dose (in terms of cell survival) in the ‘Large’ MCTS. Quantitative drug doses can be specified and implemented upon the availability of data.

### HAP and IR monotherapies attack tumours in different ways

In this initial *in silico* experiment, a MCTS is subjected to a monotherapy of either one dose of HAPs or one dose of IR. Our *in silico* results demonstrate that HAP and IR monotherapies attack the MCTS in different ways. This can be understood by regarding the treatment responses in Figure 9 and Figure 10. Figure 9 shows cell-cycle phase specific survival data, in terms of cell count over time, when the ‘Small’ or ‘Large’ MCTS is subjected to a HAP or IR monotherapy. Similarly, Figure 10 shows the composition of cells, in terms of their cell-cycle phase, in response to a HAP or IR monotherapy dose. Our results demonstrate that for the ‘Small’, well-oxygenated MCTS, HAPs have negligible effect on the cell count (see Figure 9) and, by extension, on the cell-cycle phase composition (see Figure 10). This shows that, by design, HAP treatments have little effect on tumours that are not hypoxic enough to cause significant HAP-to-AHAP bioreduction. For the ‘Large’ MCTS, however, HAPs successfully eliminate cells, particularly the inner cells of the MCTS, labeled ‘slow/non-proliferative’ (see Figure 9). This causes a change in the cell-cycle phase composition in favour of the proliferative cells in the outer shell of the MCTS (see Figure 10). Our results further show that, for both the ‘Small’ and the ‘Large’ MCTSs, IR eliminates cells of all cell-cycle states (see Figure 9), but alters the cell-cycle phase composition in favour of the inner, hypoxic cells as these are less sensitive to radiotherapy (see Figure 10). These opposing effects on the cell-cycle phase composition achieved by HAPs and IR in the ‘Large’ MCTS indicate that, for tumours that are hypoxic enough for HAPs to have an effect, HAP-IR combination treatments have the potential of producing multifaceted attacks on tumours.

**Fig 9.**
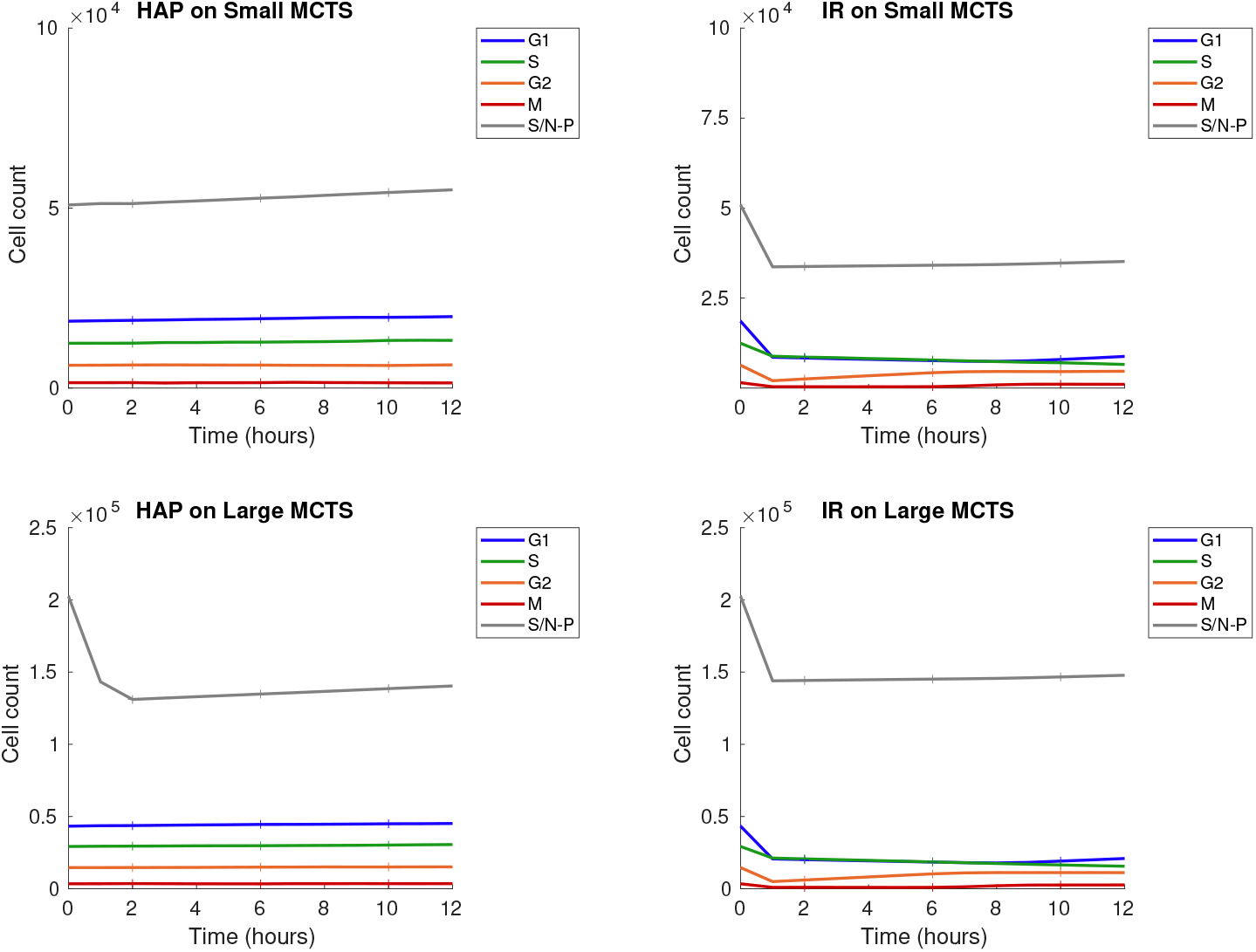
Treatment responses for HAP (left) and IR (right) monotherapies for the ‘Small’ (top) and ‘Large’ (bottom) MCTS. The monotherapy is given at *T*_0_ =0 hours. Graphs demonstrate cell-cycle specific **cell count** (i.e. number of viable, undamaged cells) over time. The slow/non-proliferative, inner cancer cells are labeled S/N-P. Solid lines show mean values, and the height of ‘+’ markers show standard deviations for 10 *in silico* runs.

**Fig 10.**
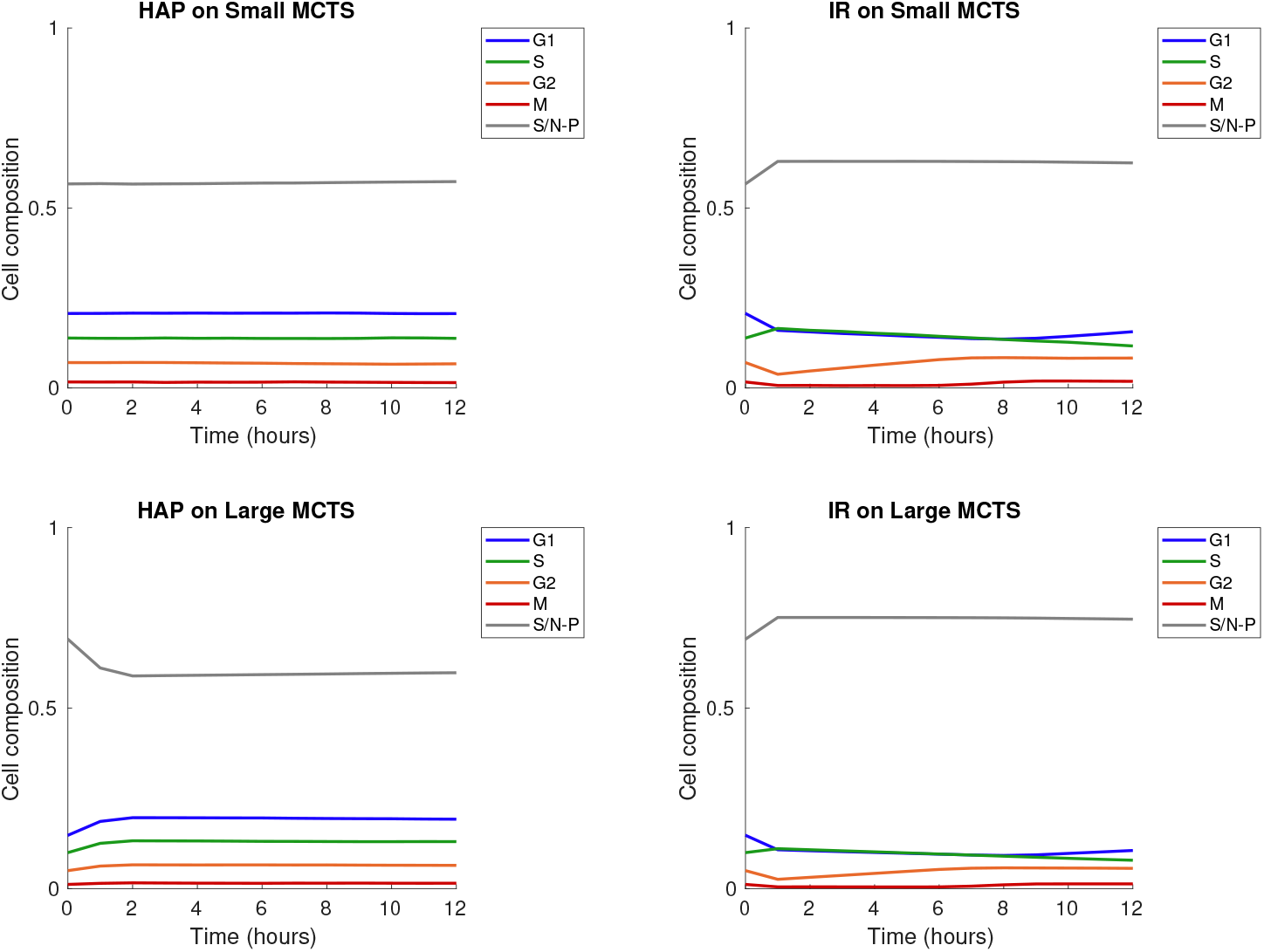
Treatment responses for HAPs (left) and IR (right) monotherapies for the ‘Small’ (top) and ‘Large’ (bottom) MCTS. The monotherapy is given at *T*_0_ = 0 hours. Graphs demonstrate cell-cycle specific **composition** (of viable, undamaged cells) over time. The slow/non-proliferative, inner cancer cells are labeled S/N-P. Solid lines show mean values for 10 *in silico* runs (standard deviations are negligible hence not shown).

Since radiation responses are enhanced by the presence of molecular oxygen, we investigated which monotherapy (i.e. HAP or IR) best eliminates hypoxic cells and re-oxygenates MCTSs after one single treatment dose. To demonstrate the overall change of oxygenation levels in the MCTSs, as a result of the monotherapies, Figure 11 provides histograms for cellular oxygenation levels at time *T*_0_ (the time of therapy administration) and at time *T*_0_ +4 hours. From this figure we can see that for the ‘Small’ MCTS, HAPs do not alter the overall intra-tumoural oxygenation but IR does, since HAPs are not efferctive but IR is. For the ‘Large’ MCTS, on the other hand, both HAPs and IR alter the overall intra-tumoural oxygenation but only HAPs manage to eliminate the most hypoxic cells, and thus shift the oxygen histogram away from the most severe levels of hypoxia. This indicates that administering HAPs as a neoadjuvant therapy prior to radiotherapy may enhance the effect of radiotherapy in tumours that are sufficiently hypoxic for HAPs to be effective.

**Fig 11.**
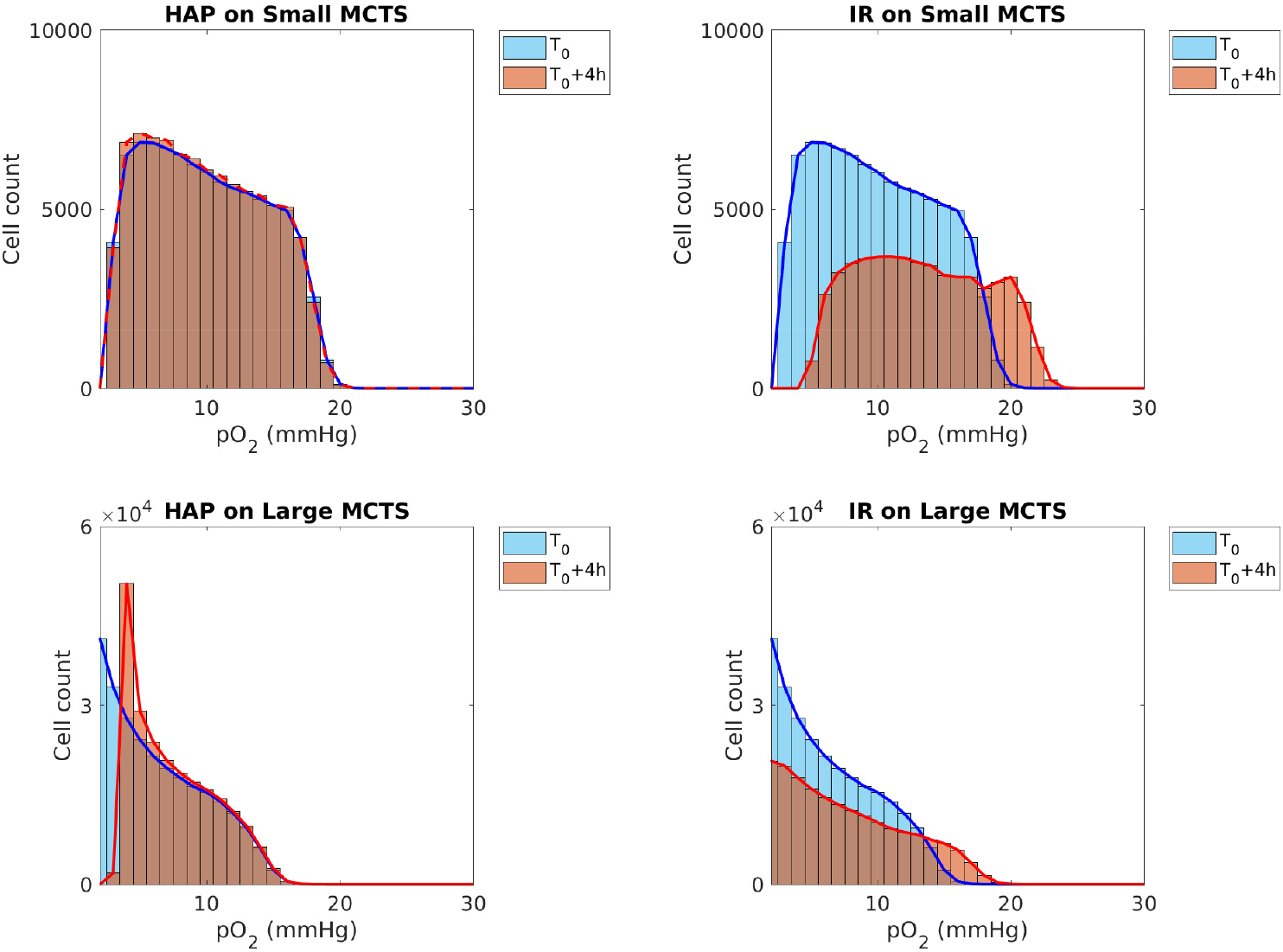
Treatment responses for HAPs (left) and IR (right) monotherapies for the ‘Small’ (top) and ‘Large’ (bottom) MCTS. Histograms over cellular oxygenation levels at time *T*_0_ (monotherapy administration time) and 4 hours later are shown. Results are based on mean values from 10 *in silico* runs.

### HAP-IR treatment scheduling impacts HAP efficacy in sufficiently hypoxic tumours

In order to study the optimal treatment scheduling of HAP-IR combination therapies, simulated MCTSs are here given one dose of HAPs and one dose of IR, using different schedules. Figure 12 shows the cell count over time when one dose of HAPs and one dose of IR are administered with various schedules. Specifically, either HAPs are given at 0 hours (followed by IR at 0, 12, 24 or 48 hours) or IR is given at 0 hours (followed by HAPs at 12, 24 or 48 hours). The results in Figure 12 demonstrate that for the ‘Small’ MCTS, scheduling does not impact the overall treatment outcome, as HAPs with the chosen [*pO*_2_]_50_ value are not effective. For the ‘Large’ MCTS however, it is here more effective to give HAPs before IR, than to give IR before HAPs. This indicates that, in tumours that are hypoxic enough for HAPs (with certain [*pO*_2_]_50_ values) to be effective, the HAP-IR treatment scheduling impacts the efficacy of the combination treatment. Note that, as is demonstrated in the supplementary material, the [*pO*_2_]_50_ value will affect the impact and importance of treatment scheduling.

**Fig 12.**
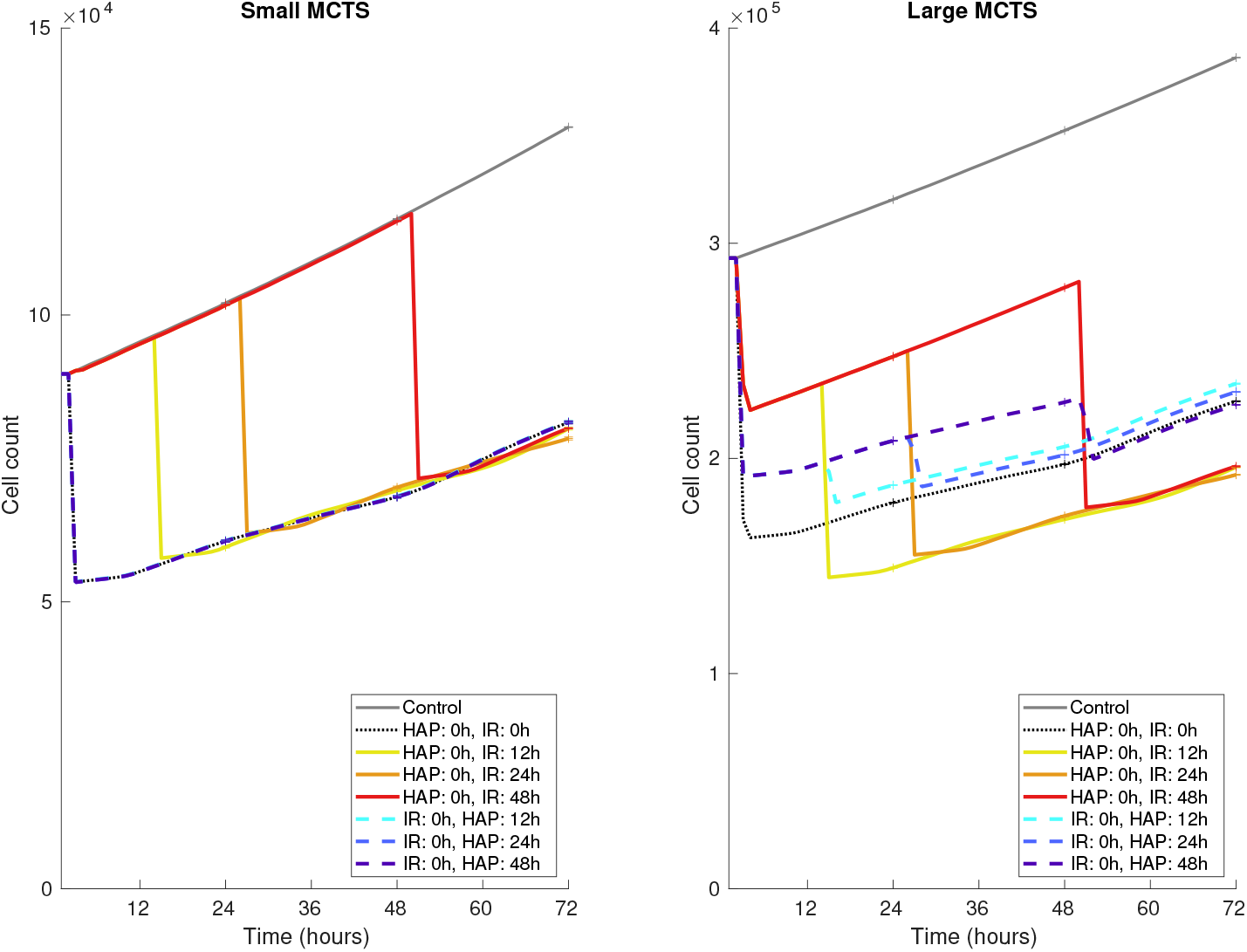
Treatment responses (in terms of cell count) for HAP-IR combination therapies in the ‘Small’ MCTS (left) and the ‘Large’ MCTS (right). One dose of HAPs and one dose of IR are administered at various schedules. Solid and dashed lines show mean values, and the height of the ‘+’ markers show standard deviations for 10 *in silico* runs.

### HAPs enhance radiotherapy effects in sufficiently hypoxic tumours

To investigate if and when HAPs enhance the effect of radiotherapy, simulated MCTSs are subjected to either IR monotherapies or HAP-IR combination therapies. In the combination therapy case, HAPs are administered at time *T*_0_ and IR is administered at time *T*_0_ + 48 hours. In the monotherapy case, radiotherapy is administered at time *T*_0_ + 48 hours. For a thorough investigation, the oxygen-levels of the ‘Large’ and ‘Small’ tumours are further scaled by multiplication with a factor 1, 1/2 or 1/4 so that we have 6 different tumours on which to test if neoadjuvant HAPs enhances radiotherapy efficacy. Figure 13 shows IR treatment responses in form of survival data (both in terms of number of surviving cells and fraction of surviving cells). From these plots we see that for very hypoxic MCTSs, the administration of neoadjuvant HAPs does increase the effect of radiotherapy. However, for well-oxygenated MCTS, neoadjuvant HAPs do not increase the effect of radiotherapy.

**Fig 13.**
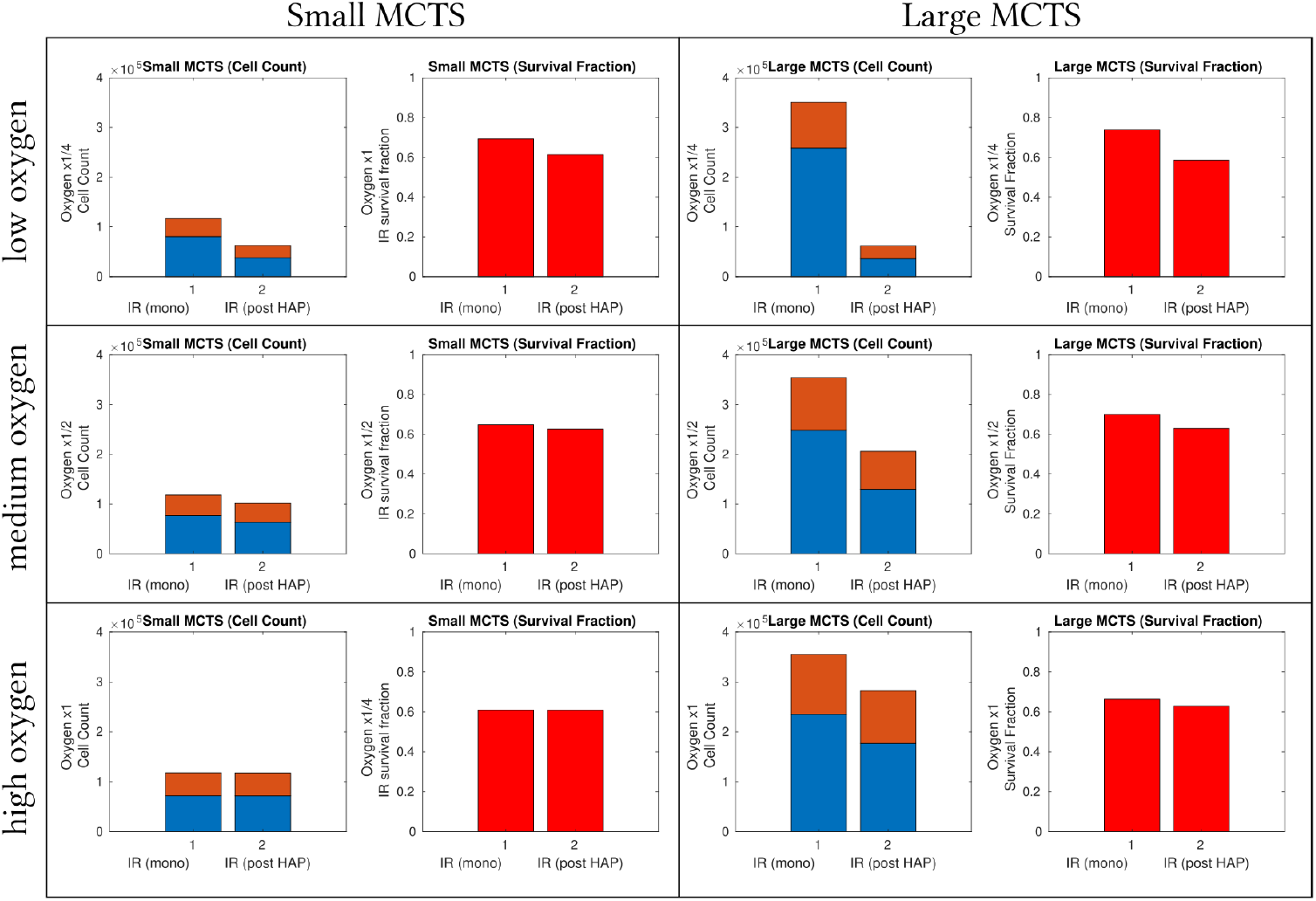
Treatment responses of radiotherapy in various MCTSs when either (1) an IR monotherapy dose is administered at *T*_0_ +48 hours or (2) IR is given at *T*_0_+48 hours following a prior HAP dose at time *T*_0_. Note that only explicit IR responses (not HAP responses) are shown. The oxygen-levels of the ‘Small’ (left) and ‘Large’ (right) tumours are scaled by a factor of 1 (least hypoxic), 1/2 or 1/4 (most hypoxic). The value calibrated from *in vitro* experiments [35] correspond to a scaling with factor 1. Orange + blue bars show number of viable cells (instantaneously) before IR administration, blue bars show the number of viable cells (instantaneously) post IR. Red bars show how many cells (as a fraction) survived the IR attack.

### The intra-tumoural oxygen landscape impacts HAP efficacy

Above, we have demonstrated various ways that the intra-tumoral oxygenation level impacts HAP and IR monotherapies and combination therapies. Further, in order to investigate if the spatio-temporal intra-tumoral oxygen landscape impacts HAP efficacy, two MCTSs with different oxygen landscapes are here compared. Omitting details of oxygen dynamics and vessel structure, hypoxic regions are here manually assigned in the MCTSs so that every cancer cell is set to be either severely hypoxic (*p*_*O*2_ = 1 mmHg) or very well-oxygenated (*p*_*O*2_ = 100 mmHg). Both MCTSs, named MCTS A and MCTS B, are assigned the same number of severely hypoxic and well-oxygenated cancer cells at the time-point when treatment commences. In MCTS A, the hypoxic region is made up of one concentric sphere in the core of the MCTS, whilst in MCTS B, the hypoxic regions consist of multiple spheres, evenly spread out across the MCTS. MCTS A and MCTS B are illustrated in Figure 14. The severely hypoxic cancer cells are here called *activator cells*, as the prodrug bioreduction (or activation) is maximal in severly hypoxic environments. The well-oxygenated cells are here referred to as *bystander cells*, as the bioreduction is minimal in well-oxygenated environments. Thus any lethal AHAP concentration occurring in a bystander cell is a result of HAP-to-AHAP bioreduction occurring outside the bystander cells.

**Fig 14.**
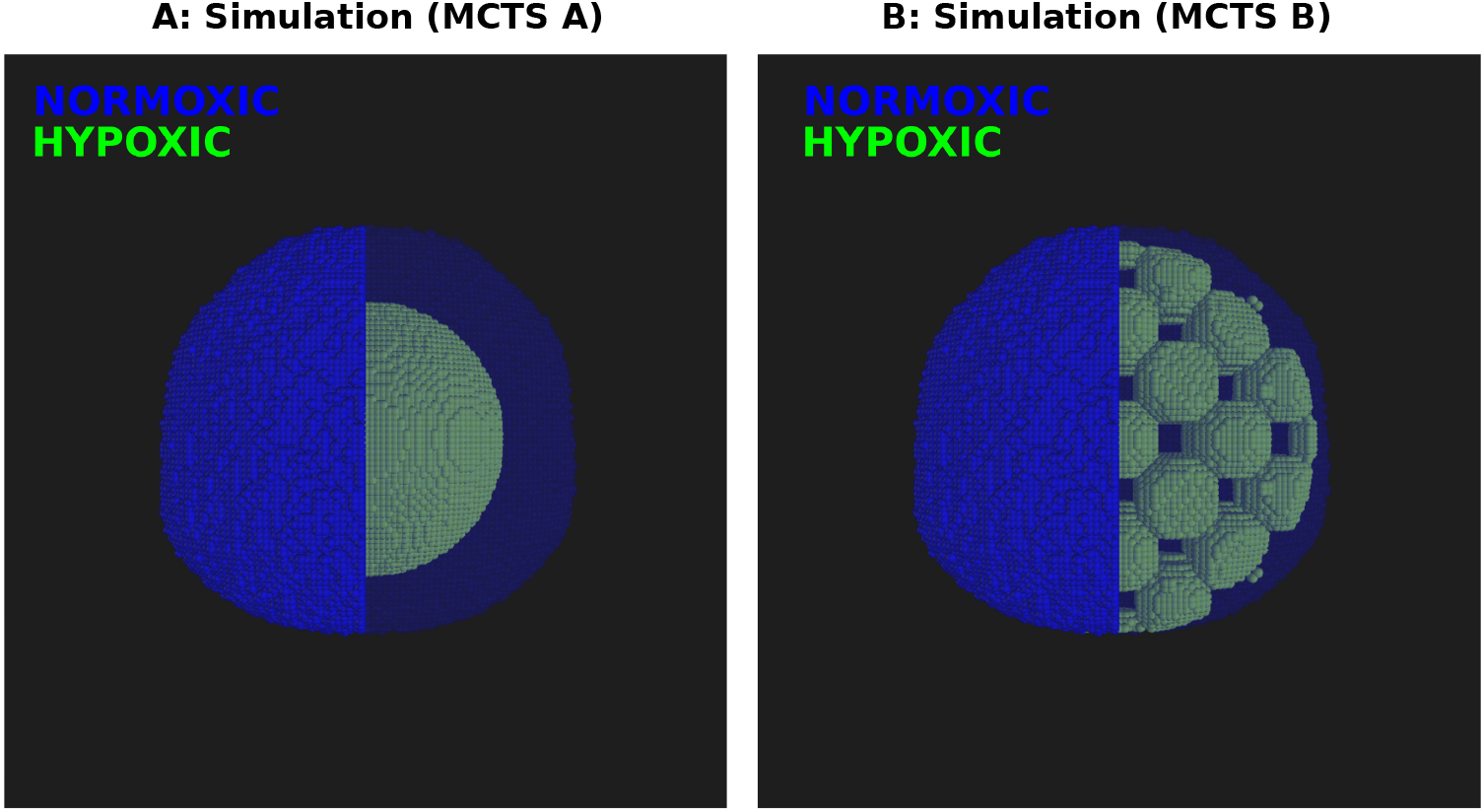
MCTS A and B prior to treatment commencing. The MCTSs are visualised in both opaque and transparent formats. Hypoxic activator cells are shown in green and normoxic bystander cells are shown in blue. Activator and bystander cells are manually set so that MCTSs A and B contain the same number of activator and bystander cells before treatment commences.

From Figure 15 it is clear that the bystander effects are higher in MCTS B than in MCTS A, although all activator cells are eliminated in both MCTSs. When the activator cells are spread out across the spheroid, as in MCTS B, there are more interfaces in which bystander cells experience significant bystander effects. Although the oxygen landscape in MCTS B is highly synthetic, this *in silico* experiment shows that the intra-tumoural oxygen landscape does impact the efficacy of HAPs. In the supplementary material, more MCTSs with distinct oxygen landscapes, subjected to HAP monotherapies are explored.

**Fig 15.**
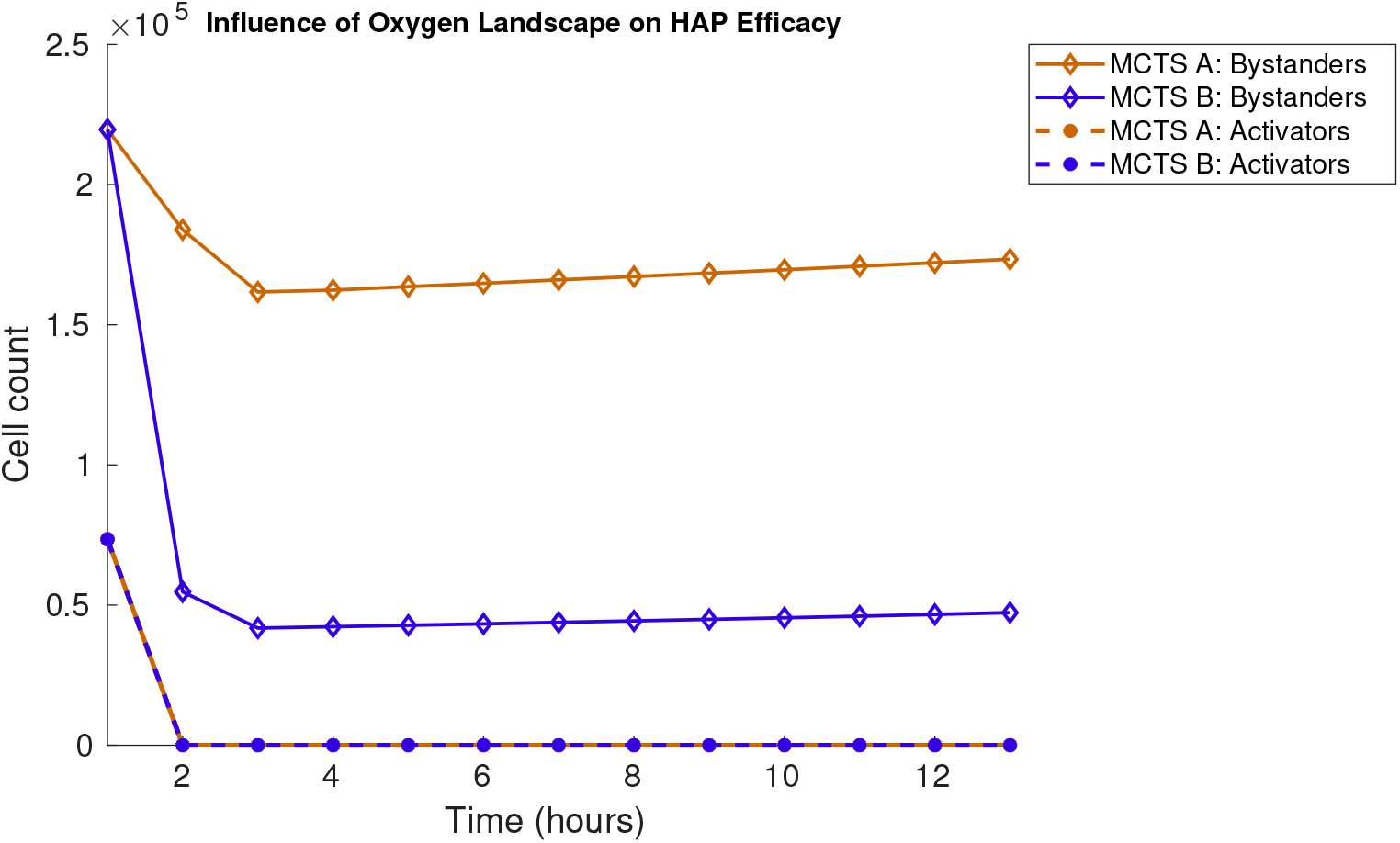
Treatment responses in MCTS A and MCTS B when HAPs are administered at 0 *T*_0_ = 0 hours. The number of viable (undamaged) cells are plotted over time for MCTS A and MCTS B. Cell counts for activator cells (pO_2_ = 1 mmHg) are shown in dashed lines and bystander cell counts (pO_2_ = 100 mmHg) are shown in solid lines. Results demonstrate mean values for 10 *in silico* runs.

## Conclusion

Previous *in vitro* and *in vivo* studies have validated the successfulness of HAPs in laboratory settings, however, this preclinical success has not yet been reflected in clinical trials. In an attempt to elucidate the unsatisfactory results from clinical HAP trials, we in this study investigate how oxygen-related tumour features and treatment scheduling impact the efficacy of HAP monotherapies and HAP-IR combination therapies *in silico*. To this end, we have developed a mathematical model capturing the spatio-temporal dynamics of tumours subjected to multimodality treatments comprising HAPs and IR. A set of key results (*i* to *iv*) relating to (Class II) HAP efficacy *in silico* have here been demonstrated.

i. HAPs and IR attack tumours in different, complementary, ways. Whilst IR provides a highly effective way to kill cancer cells, tumour regions containing hypoxic and resting cells are significantly more resistant to IR than are tumour regions with well-oxygenated and actively cycling cells. HAPs, however, are alkylating agents which bioreduce in (primarily) hypoxic areas, hence HAPs mainly inflict damage in hypoxic tumour regions. Consequently, HAP-IR combination treatments have the potential to produce multifaceted attacks on tumours with heterogeneous oxygen landscapes.
ii. HAP-IR treatment scheduling may impact treatment efficacy. The impact of scheduling is apparent in tumours that contain regions that are hypoxic enough for IR to be ineffective (when the HAP bioreduction is, in most part, restricted to occur in those regions).
iii. HAPs may function as IR treatment intensifiers in tumours that contain hypoxic regions in which IR is ineffective.
iv. Not only the overall intra-tumoural oxygenation levels, but also the intra-tumoural oxygen landscape, impacts the efficacy of HAP monotherapies.

In this study, we qualitatively investigated various aspects of HAP-IR treatment schedules using a multiscale mathematical framework. Upon the availability of *in vitro* and *in vivo* data, this mathematical framework can be calibrated in order to serve as an *in silico* testbed for predicting HAP-IR treatment scenarios. As a result of interdisciplinary collaborations, the mathematical framework used in this study has previously been validated *in vitro* and *in vivo* for applications other than HAP-IR combination treatments [37,56]. The multiscale nature of the framework enables integration of data from various scales, be it from the subcellular scale, the cellular scale or the tissue scale. As an example of useful data, the multi cellular tumour spheroid data previously produced by Voissiere *et al.* [35] provided our framework with calibration data for tumour growth and spatio-temporal oxygen dynamics. Using existing experimental data to create data-driven mathematical models is a resourceful step involved in the advancement of mathematical oncology [57].

In a recent publication, Spiegelberg *et al.* [19], claim that the (lack of) clinical progress with HAP-treatments can, in great part, be attributed to the omission of hypoxia-based patient selection. Our *in silico* study demonstrates that whilst (class II) HAPs are effective treatment intensifiers for sufficiently hypoxic tumours, they have negligible effect on more well-oxygenated tumours. In simple terms: some tumours are suitable to be paired with treatment plans involving HAPs whilst others are not. In line with Spiegelberg *et al.*’s claims [19], our *in silico* results indicate that a personalised medicine approach is preferable if treatments involving HAPs (that are similar to TH-302) are to achieve their maximum potential in clinical settings, where intra-tumoural oxygenation status can be assessed in multiple ways: By inserting oxygen electrodes into tumours, *pO*_2_ values can directly be measured, but this measuring technique is invasive and does not distinguish between hypoxic and necrotic tumour regions [19]. Alternatively, less invasive imaging techniques, such as positron emission (PET-scans) and oxygen-enhanced magnetic resonance (MRIs), can be used to evaluate oxygen levels in tumours [2,19]. Moreover, there now exist several hypoxia gene expression signatures that may be used to characterise hypoxia-related tumour features, and some of these signatures have been conferred with poor clinical prognoses [19]. Avoiding a tumour biopsy, by measuring hypoxia secreted markers in the blood, would, furthermore, constitute a more expeditious way to assess tumour hypoxia [19]. Without further discussing the advantages and disadvantages of various hypoxia assessment methods, the above discussion illustrates that it is, indeed, feasible to invoke stricter selection regimes when deciding whether or not to pair tumours with HAP treatments in clinical trials [19], in line with a personalised medicine approach.

## Acknowledgements

SH was supported by the Medical Research Council [grant code MR/R017506/1] and Swansea University PhD Research Studentship. MK acknowledges the financial support from the Canadian Institutes of Health Research (CIHR). LJD, AY and PL acknowledge financial support from ERC advanced grant (ERC-ADG-2015, n° 694812 – Hypoximmuno) and EUROSTARS (COMPACT 12053).

## Supplementary Material

**SM1a:** Complement to Figure 12 – *HAP-IR treatment scheduling impacts HAP efficacy in sufficiently hypoxic tumours*.

**SM1b:** Complement to Figure 12 – *The* [*pO*_2_]_50_ *value influences scheduling outcomes*.

**SM2:** Complement to Figure 13 – *HAPs enhance radiotherapy effects in sufficiently hypoxic tumours*.

**SM3:** Complement to Figure 15 – *The intra-tumoural oxygen landscape impacts HAP efficacy*.

**SM4:** *Pseudo-code flowchart*.

### SM1a Complement to Figure 12

Figures 16 and 17 show that the scheduling-experiment, with results provided in Figure 12 in the main manuscript, are qualitatively the same if a damaged cell is instantly removed from the lattice (Figure 16) or if a damaged cell is moved from the lattice after a time period corresponding to its doubling time (Figure 17).

**Fig 16.**
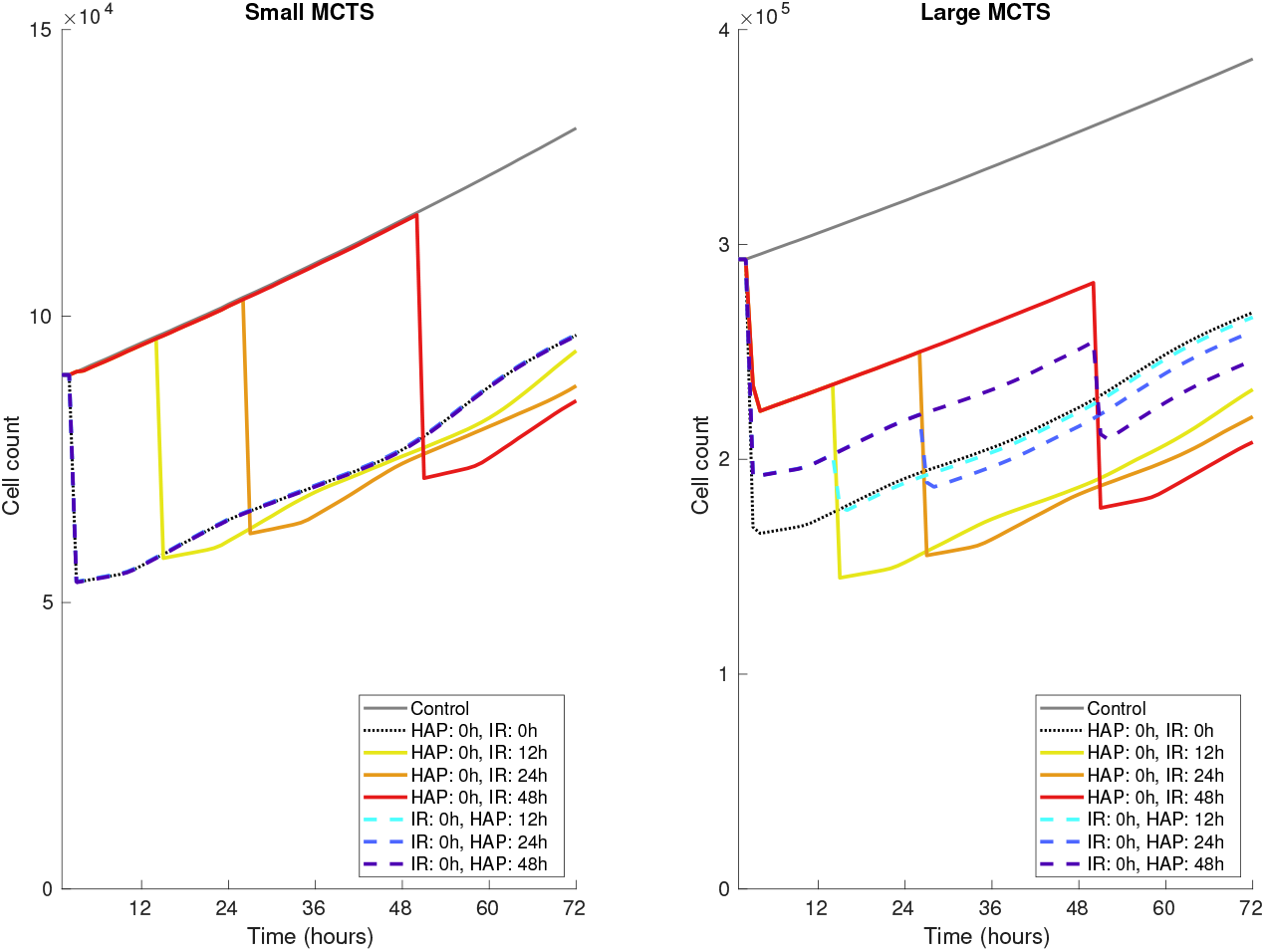
Scheduling of HAP-IR combination treatments, complement to Figure 12 in the main manuscript. Cells are removed from the lattice instantaneously after the lethal event occurred.

**Fig 17.**
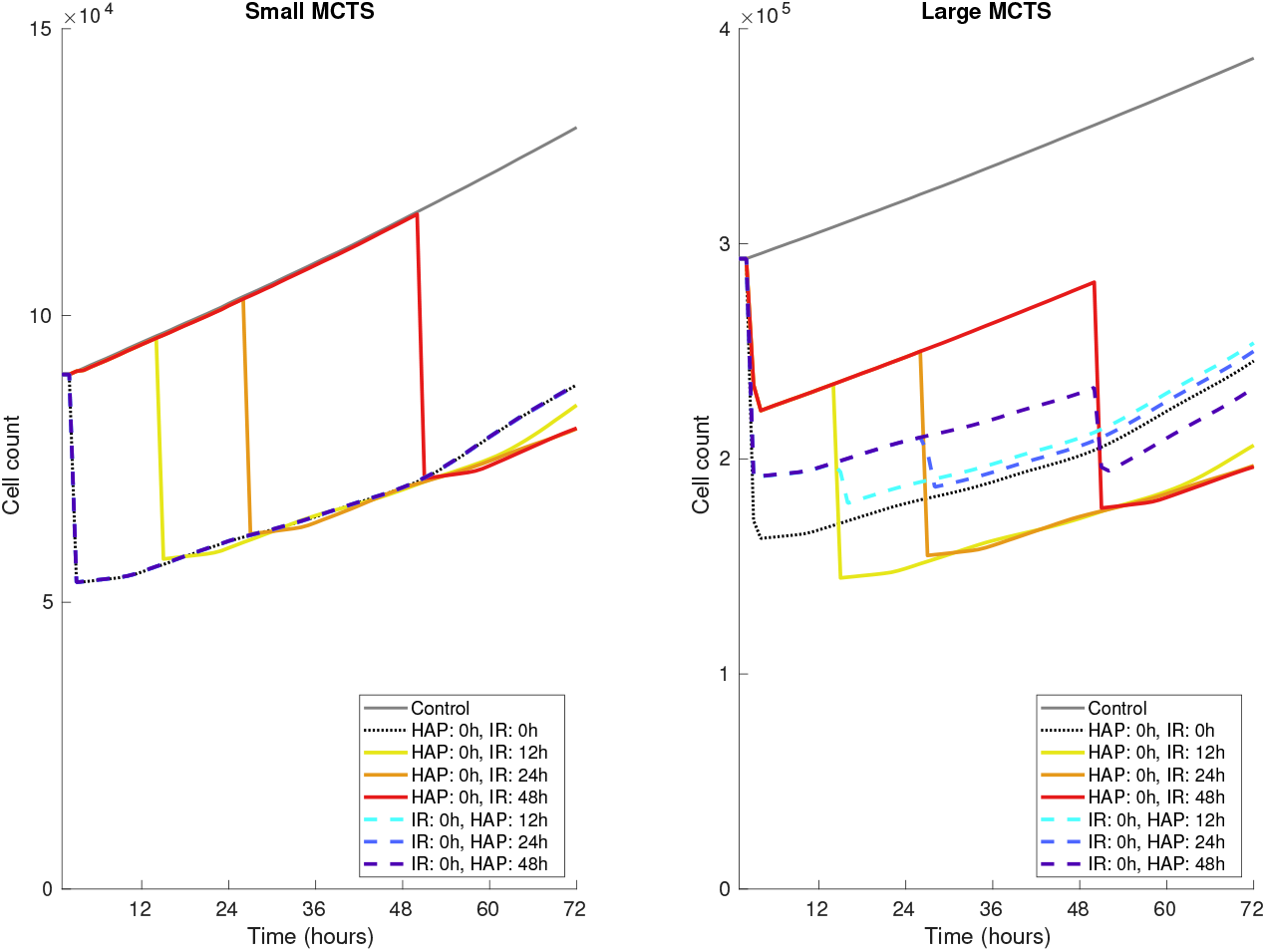
Scheduling of HAP-IR combination treatments, complement to Figure 12 in the main manuscript. Cells are removed from the lattice after a time corresponding to their doubling time (*τ_i_*) post the lethal event.

### SM1b The [*pO*_2_]_50_ value influences scheduling outcomes

The [*pO*_2_]_50_ parameter value, denoting the oxygen value yielding 50% HAP-to-AHAP hourly bioreduction (see Figure 5), impacts the efficacy of various HAP-IR combination therapy schedules. To demonstrate this, Figures 18 (left) and 18 (right) respectively show the cell count over time when the ‘Small’ and ‘Large’ tumour (illustrated in Figure 8) are subjected to various HAP-IR schedules. In Figure 18, the [*pO*_2_]_50_ value has been increased by a factor of 5 from its original value used in the *in silico* experiments described in the manuscript (see Figure 12).

**Fig 18.**
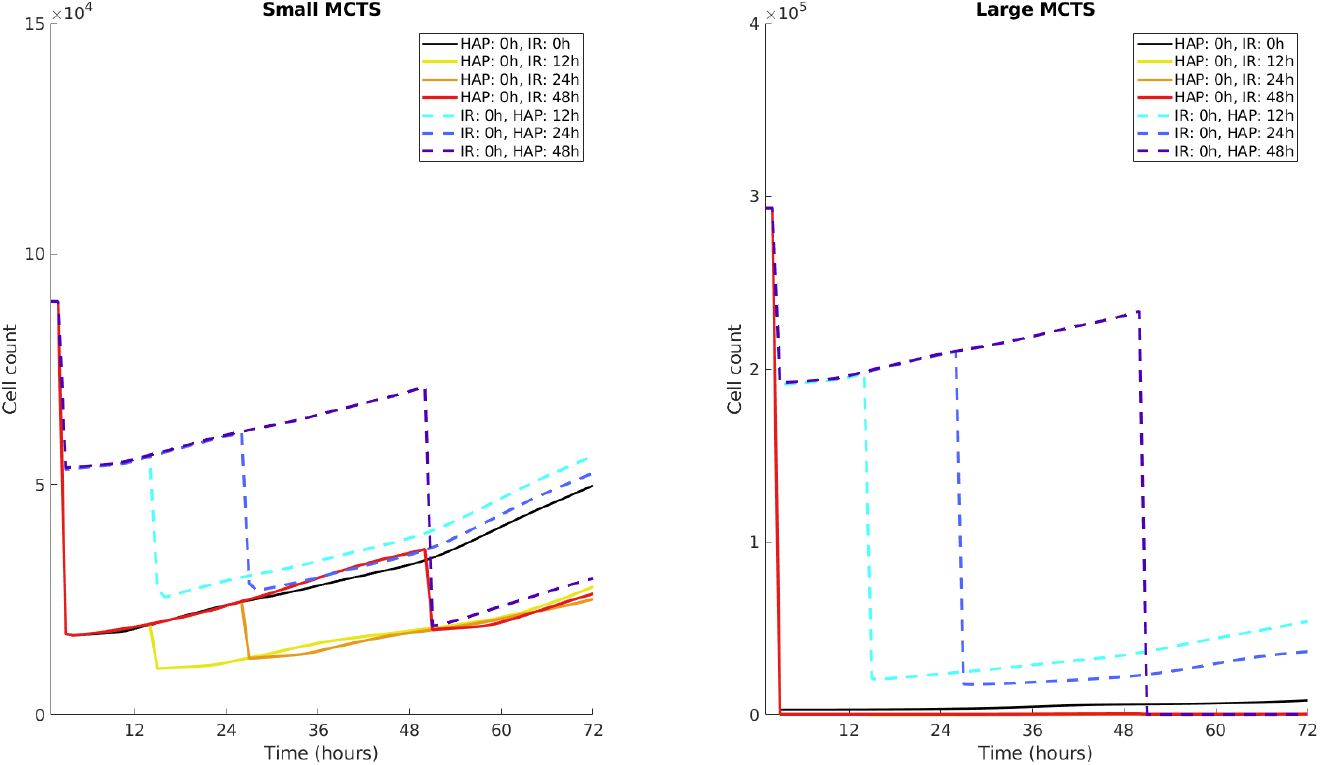
Scheduling of HAP-IR combination treatments, Complement to Figure 12. The [*pO*_2_]_50_ value is 5 times larger than in the original *in silico* experiments described in the manuscript.

### SM2 Complement to Figure 13

Figures 19 and 20 show that the experiment that investigates if HAPs act as radiotherapy enhancers, with results provided in Figure 13, are qualitatively the same if a damaged cell is instantly removed from the lattice (Figure 19) or if a damaged cell is moved from the lattice after a time period corresponding to its doubling time (Figure 20).

**Fig 19.**
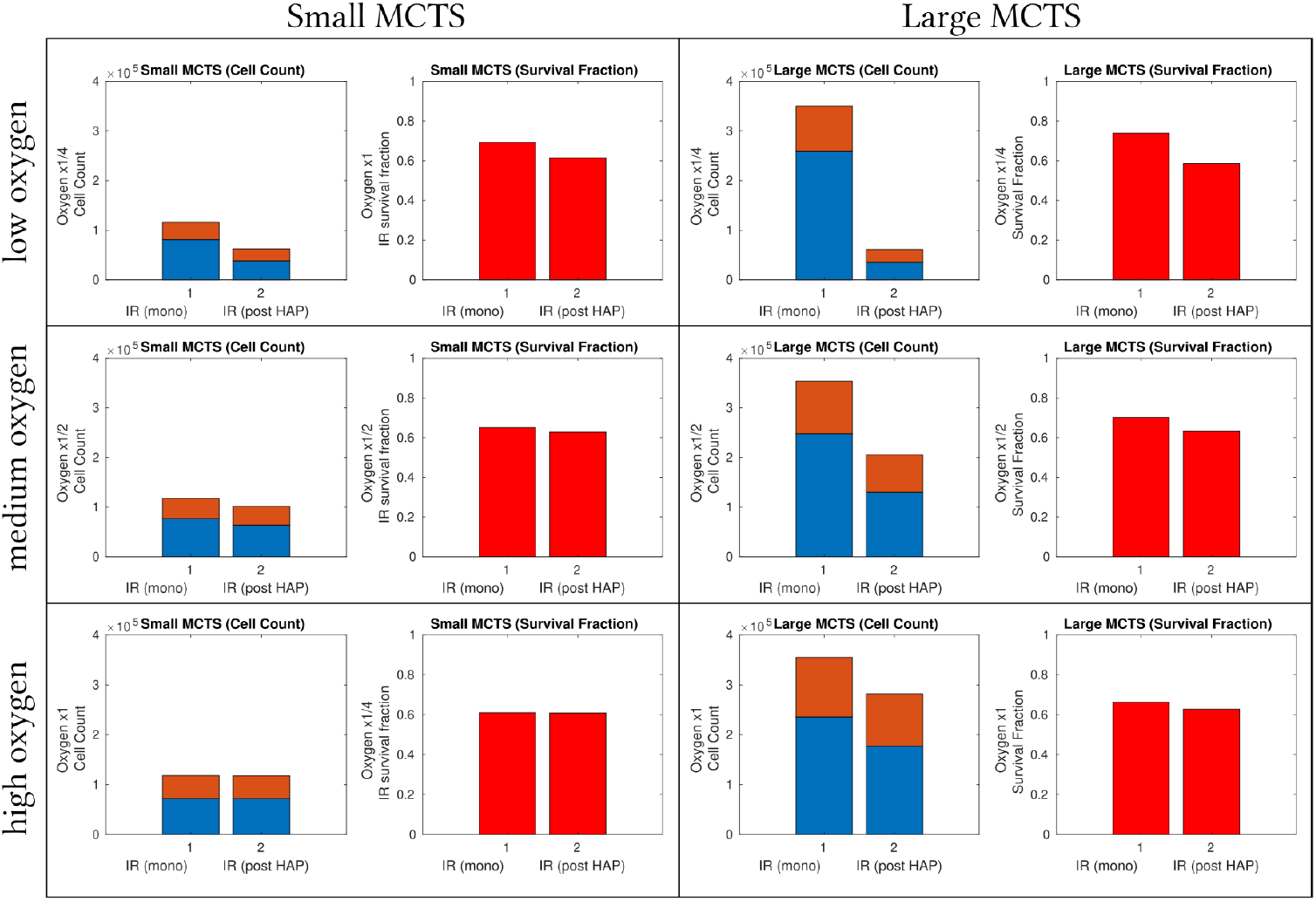
Treatment responses of radiotherapy in various MCTSs when either (1) an IR monotherapy dose is administered at *T*_0_ +48 hours or (2) IR is given at *T*_0_+48 hours following a prior HAP dose at time *T*_0_. Complement to Figure 13. Cells are removed from the lattice instantaneously after the lethal event occurred.

**Fig 20.**
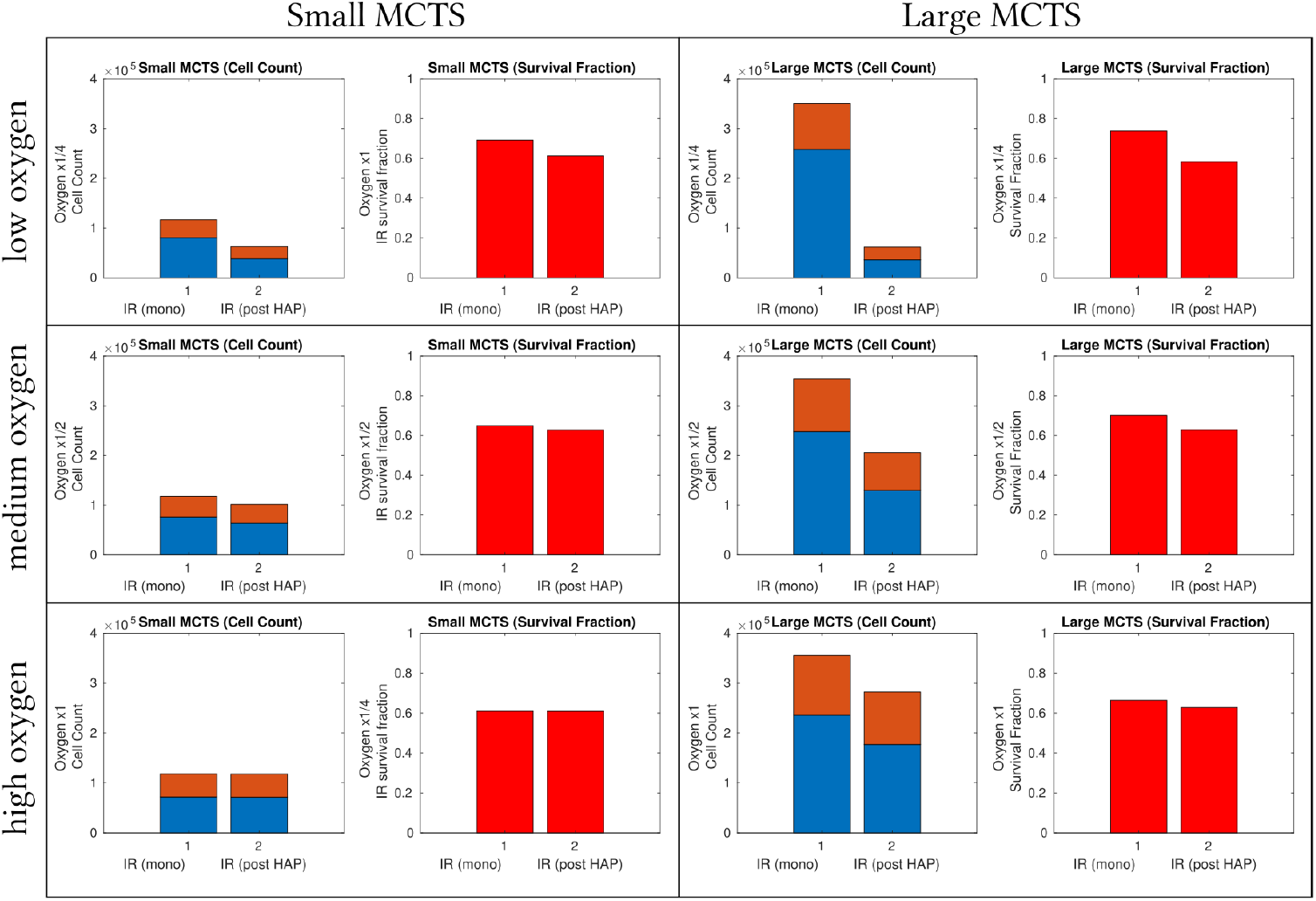
Treatment responses of radiotherapy in various MCTSs when either (1) an IR monotherapy dose is administered at *T*_0_ +48 hours or (2) IR is given at *T*_0_+48 hours following a prior HAP dose at time *T*_0_. Complement to Figure 13. Cells are removed from the lattice after a time corresponding to their doubling time (*τ_i_*) post the lethal event occurred.

### SM3 Complement to Figure 15

As a complement to Figure 15, and the investigation concerning how the intra-tumoural oxygen landscape impacts HAP efficacy, we here introduce three more *in silico* tumour spheroids: MCTS C, D and E, in addition to MCTS A and B introduced in Figure 14 in the manuscript. The MCTSs are visualised in Figure 21, where all MCTSs contain the same number of hypoxic (oxygen level: 1 mmHg) and well-oxygenated (oxygen level: 100 mmHg) cells before treatment commences. The cell count over time when each of the MCTSs are subjected to a HAP dose at zero hours is available in Figure 22, which illustrates that the oxygen landscape impacts how many cells survive the HAP treatment.

**Fig 21.**
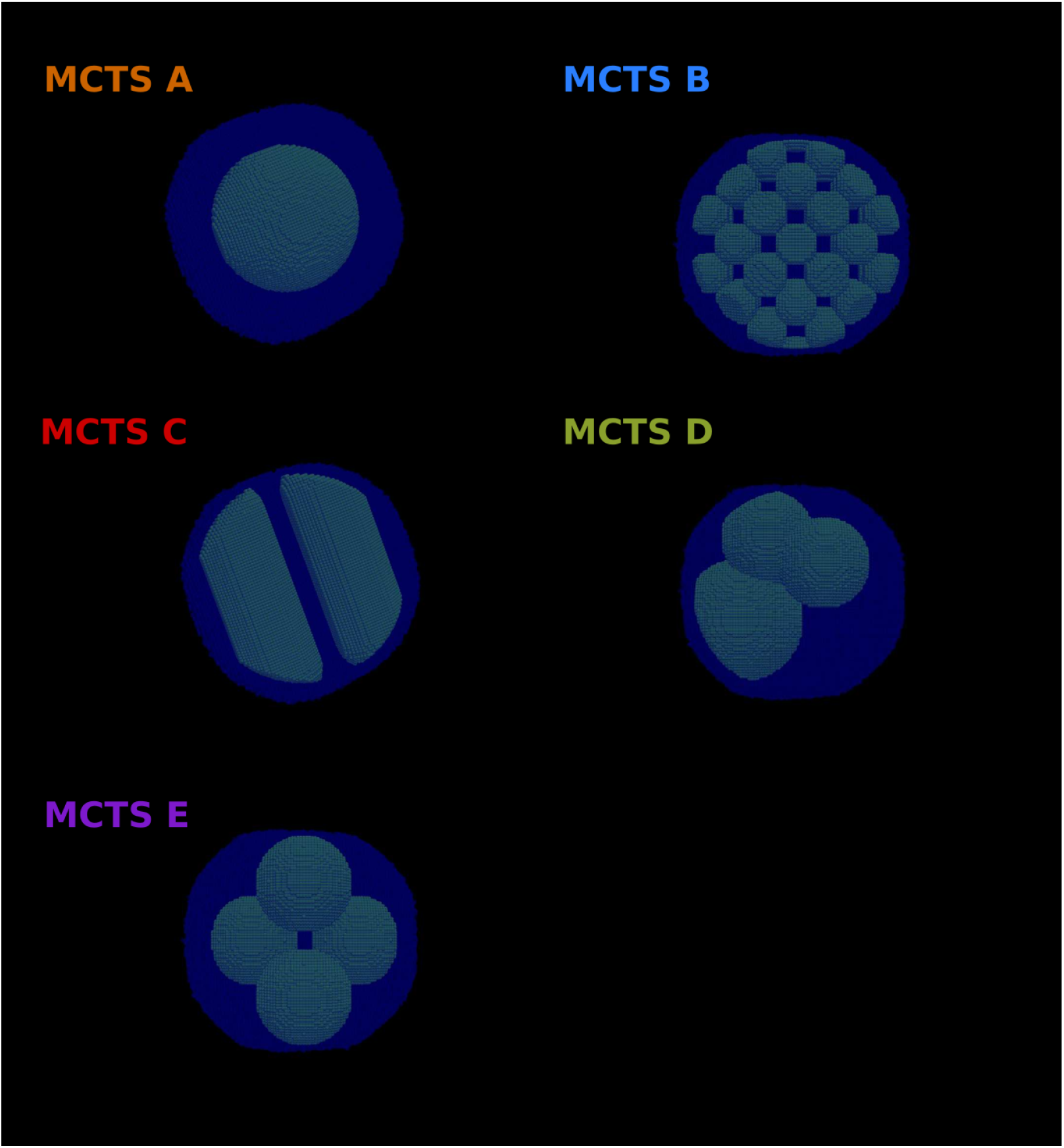
MCTSs A-E all comprise the same number of hypoxic and well-oxygenated cells, but the hypoxic cells are clustered in different ways in the various MCTSs. Green cells are hypoxic (oxygen level: 1 mmHg) and blue cells are well-oxygenated (oxygen level: 100 mmHg).

**Fig 22.**
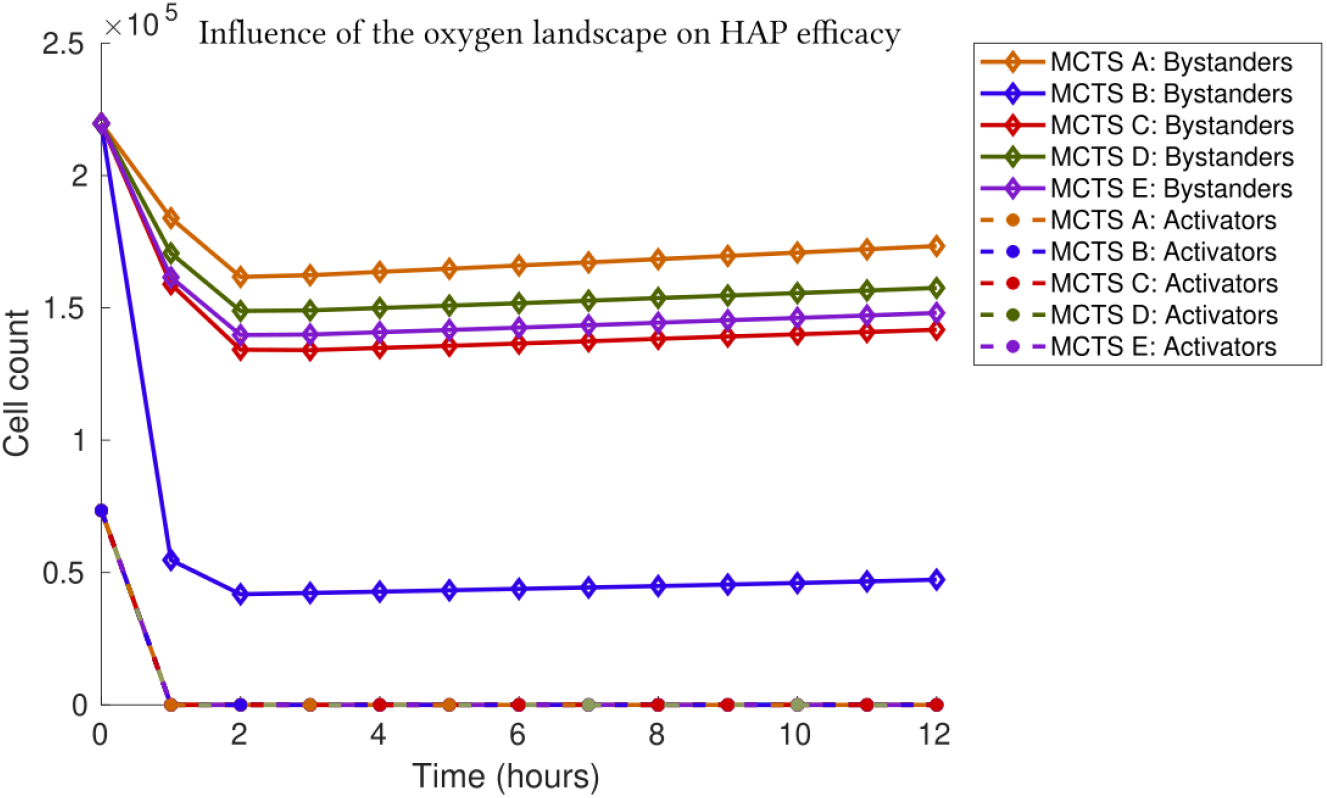
Cell count over time when MCTSs A-E are subjected to a HAP dose at zero (0) hours. Mean values, based on 10 *in silico* runs, are shown. The resulting standard deviations are less than 0.5% of the means and hence not visible in the plot.

### SM4 Pseudo-code flowchart

A diagrammatic representation of the code used in this study is provided in Figure 23.

**Fig 23.**
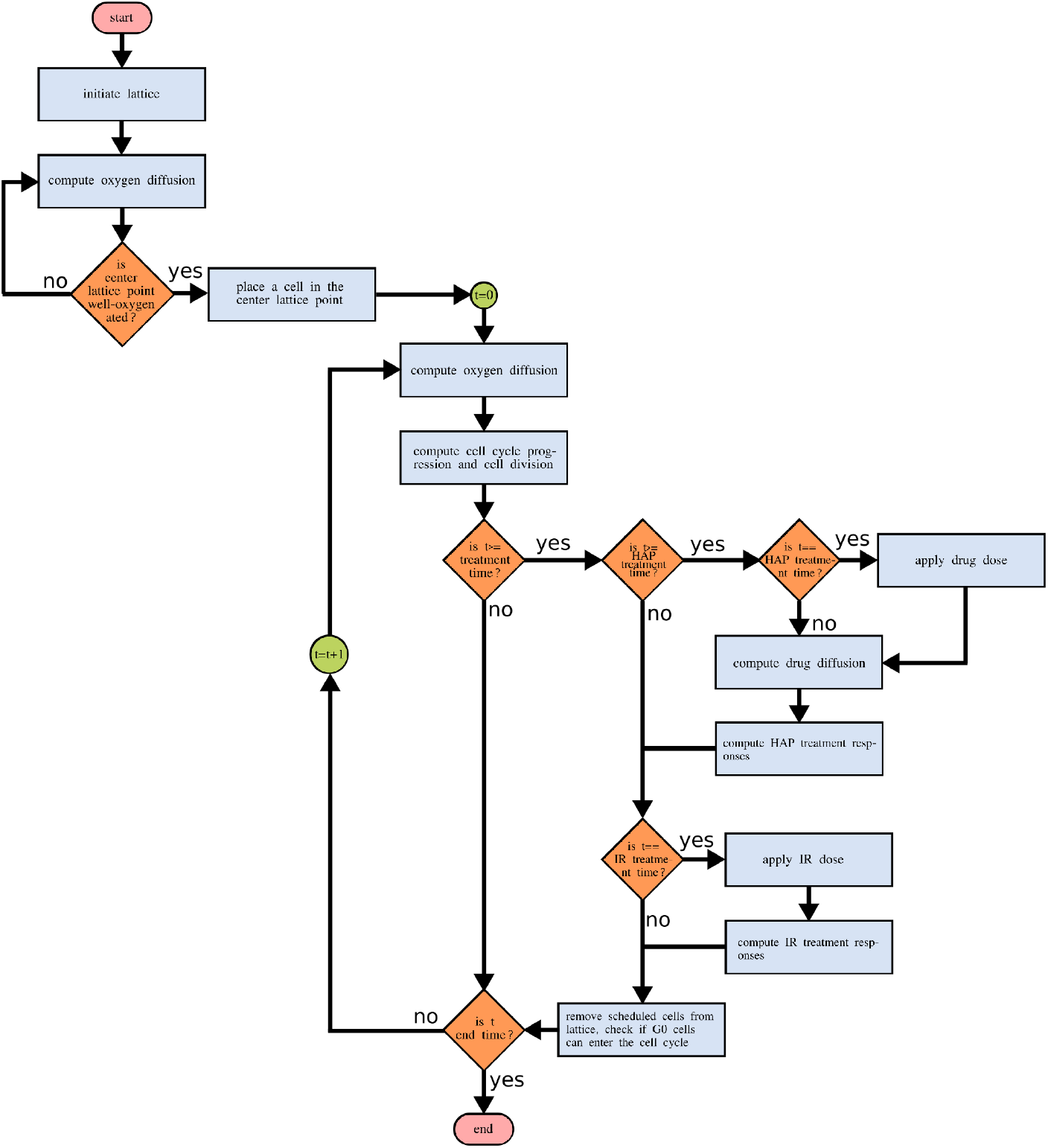
A pseudo-code flowchart describing the basic structure of the in *silico* experiments. An in-house C++ code is used for model implementation.

